# Striatal neurons are recruited dynamically into collective representations of self-initiated and learned actions in freely-moving mice

**DOI:** 10.1101/2022.10.30.514405

**Authors:** Lior Tiroshi, Yara Atamna, Naomi Gilin, Joshua A. Goldberg

## Abstract

Striatal spiny projection neurons are hyperpolarized-at-rest (HaR) and are driven to spike threshold by a small number of powerful inputs – an input-output configuration that is detrimental to response reliability. Because the striatum is important for habitual behaviors and goal-directed learning, we conducted microendoscopic imaging in freely-moving mice that express a genetically-encoded calcium indicator sparsely in striatal HaR neurons to compare their response reliability during self-initiated movements and operant conditioning. The sparse expression was critical for longitudinal studies of response reliability, and for studying correlations among HaR neurons while minimizing spurious correlations arising from contamination by the background neuropil signal. We found that HaR neurons are recruited dynamically into action representation through a moment-by-moment formation of distinct cell assemblies. While individual neurons respond with little reliability, the population response remained reliable. Moreover, we found evidence for the formation of correlated cell assemblies during conditioned (but not innate) behaviors, but this correlation was independent of the distance between neurons.

## Introduction

A critical role of the central nervous system (CNS) is to maintain a constant, robust, coherent and reliable representation of the external world and internal state of the organism. Individual sensory neurons and motoneurons need to represent their inputs and output (respectively) both reliably and reproducibly (Jones et al., 2004; Slater, 2008). However, deep within the CNS where the representations are presumably distributed within neuronal networks (perhaps with redundancy), the reliability of an individual neuron’s responses may be less important (Benedetti et al., 2009; Gur and Snodderly, 2006; Kara et al., 2000; Shadlen and Newsome, 1998). Nevertheless, the degree of reliability of neuronal responses within a given brain region will determine the resources required to represent the external world and encode actions (Bullock, 1970; Nolte et al., 2019; Shadlen and Newsome, 1998). Response reliability also has significant implications for the nature of the neural code: with reliable responses, fewer neurons are required to achieve a robust representation (Gallant and Vinje, 2001; Mainen and Seinowski, 1995).

If we consider the relationship between the size of the neuron’s individual synaptic inputs [e.g., excitatory post-synaptic potentials (EPSPs)] (Gutkin et al., 2003; Hunter and Milton, 2003) and the depolarization required to reach action potential (AP) threshold (Naundorf et al., 2006; Puelma Touzel and Wolf, 2015; Schreiber et al., 2009; Tchumatchenko et al., 2011), we can come by, even within the same brain regions, various scenarios. In one scenario, the neuron’s membrane potential is close to AP threshold (or the neuron may be a pacemaker that approaches AP threshold autonomously), and individual EPSPs are typically weak (Naundorf et al., 2006; Puelma Touzel and Wolf, 2015; Tchumatchenko et al., 2011). In this case, an external event or action is typically encoded by many (e.g., even thousands of) afferent EPSPs, such that the noisy nature of individual EPSPs will be averaged out, creating a relatively high-fidelity and reproducible representation of the input. Such a neuron would be expected to respond reliably to this representation. In another scenario, a neuron has a very hyperpolarized resting membrane potential but receives larger afferent EPSPs. In this case, because the summation of fewer inputs is sufficient to reach AP threshold, there will be less averaging out of the input variability. Thus, the responding neuron is likely to represent the event or action less reliably and less reproducibly.

In the striatum – a major basal ganglia (BG) input nucleus, integrating information from a wide range of brain regions (Doig et al., 2010; Flaherty and Graybiel, 1993; Hunnicutt et al., 2016; McFarland and Haber, 2000; Smith et al., 2004; Yeterian and Van Hoesen, 1978) – both of these scenarios can be found. The primary inputs to the striatum are excitatory projections from the cortex and thalamus, with inputs from various cortical and thalamic sub-regions converging onto the same striatal neurons (Inase et al., 1996; McFarland and Haber, 2000; Takada et al., 1998; Takadá et al., 1998). Cholinergic interneurons (CINs) and somatostatin positive low-threshold spiking interneurons (LTSIs), two spontaneously active striatal populations (Bennett et al., 2000; Tepper and Koós, 2016) – whose voltage trajectory is driven autonomously to AP threshold – receive weak cortical and thalamic inputs (Johansson and Silberberg, 2020). In contrast, spiny projection neurons (SPNs), the majority of striatal neurons (Gerfen, 1988; Graveland and Difiglia, 1985; Tepper et al., 2008), and parvalbumin positive (PV) interneurons that mediate powerful feedforward inhibition and shape striatal output through the innervation of SPNs (Assous and Tepper, 2018) are both hyperpolarized-at-rest (HaR),, and receive stronger inputs from these structures (Johansson and Silberberg, 2020). Therefore, the striatum is a good testing ground for the relationship between how individual neurons integrate their inputs and the degree of response reliability they exhibit: the autonomous pacemakers (CINs and LTSIs) should exhibit a higher degree and the HaR neurons (SPNs and PV interneurons) should exhibit a lower degree of reliability in the representation of stimuli and actions. Indeed, the tonically active neurons (TANs) of the striatum, comprised mostly of CINs (Aosaki et al., 1995), exhibit a stereotypic burst response to salient inputs, and when comparing the average response (e.g., their peristimulus time histogram) to the individual trials, it is clear that not only do individual trials reliably replicate the mean population response (Apicella et al., 1997; Kimura et al., 1984; Raz et al., 1996), the vast majority (50-90%) of TANs respond to the salient input (Apicella et al., 1997; Kimura et al., 1984; Raz et al., 1996). Thus, autonomously active CINs indeed respond robustly reliably and reproducibly to behavioral events.

In the current study, we quantified the degree of reliability of the neural responses of HaR striatal neurons that typically receive few strong individual inputs in freely-moving mice. While we predict that the overall reliability will be low, we chose to compare it between two types of naturalistic behaviors: self-initiated movement vs. operant conditioning. We hypothesized that the degree of reliability is higher for conditioned responses whose behavioral value is presumably higher than innate behaviors. To this end, we preformed microendoscopic Ca^2+^ imaging in a transgenic mouse strain that expresses the Ca^2+^ indicator GCaMP6f (Chen et al., 2013) in a sparse population comprised almost exclusively by SPNs and PV interneurons – two HaR striatal subtypes that receive strong cortical and thalamic innervation (Johansson and Silberberg, 2020). This sparse expression enabled us to clearly identify and compare the responses of the same individual neurons across sessions, days, and even weeks apart, assessing their response reliability over time and across behavioral states.

In addition to quantifying the reliability of individual neurons, we leveraged our ability to simultaneously image the activity of dozens of isolated striatal neurons to explore the correlations between their signals. We hypothesized that the low reliability of individual neurons is compensated by the on-the-fly formation of co-active clusters of neurons around a given action or stimulus. Recent studies have shown that SPNs form spatially compact neural clusters, in which nearby neurons are co-active and code for the same type of movement (Barbera et al., 2016; Parker et al., 2018; Shin et al., 2020). Importantly, the clusters in these previous studies were extracted from groups of thousands of simultaneously imaged neurons. However, the endoscopic imaging of densely packed neurons gives rise, in addition to the somatic signals, to an extensive background signal arising from neuropil activity. Recent studies have characterized these neuropil signals and concluded that they are highly correlated in space and represent a collective neural activity distinct from that of the somata (Legaria et al., 2021; Rehani et al., 2019). It is thus possible that the spatially correlated neuropil signal introduces spurious correlations among SPNs, creating a semblance of compact cell clusters operating together. Focusing on a sparse striatal population of SPNs allows for a clear separation between sources, and could help resolve the question of distance-dependent correlations among SPNs in a behaving mouse.

## Results

### Self-initiated movement in freely moving mice strongly modulates the activity of HaR striatal neurons

To examine the response reliability of striatal neurons (presumably to cortico- and thalamostriatal inputs) during naturalistic behaviors, we used sparse GCaMP mice expressing the genetically encoded calcium indicator (GECI) GCaMP6f in a sparse population of striatal neurons consisting of a majority of ~60% SPNs and ~30% PV interneurons (see Methods, Fig. S1) – two striatal populations that are HaR (Hammond, 2015; Koós and Tepper, 2002; Tepper and Koós, 2016) and typically driven to spike by few large synaptic inputs (Johansson and Silberberg, 2020). Mice implanted with a GRIN lens and microendoscopes were used to simultaneously visualize multiple sparsely distributed neurons in the dorsal striatum of 7 freely moving mice during innate behavior (Fig. 1A-C). The kinematics of the mice were monitored using video cameras, an accelerometer, and tracking software that were synchronized with the Ca^2+^ imaging, enabling us to compare the collective neuronal activity in different behavioral states.

**Figure 1.**
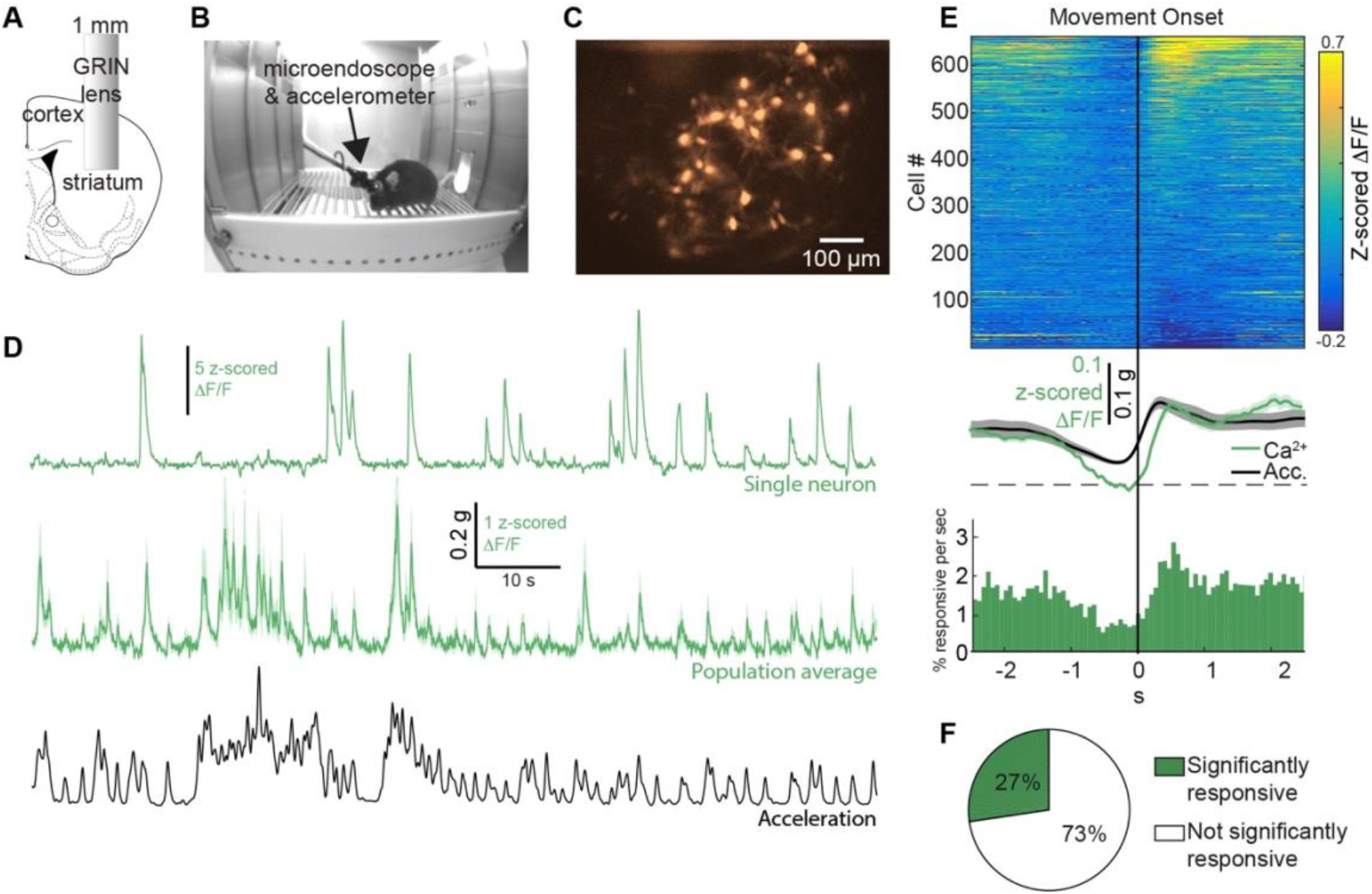
The collective neuronal activity is strongly modulated by self-initiated movement in freely moving mice. **A:** A 1-mm-diameter GRIN lens is implanted into the dorsolateral striatum. **B:** Implanted mouse with a microendoscope and an accelerometer mounted on its head moves freely in a behavior chamber. **C:** Image via lens in freely moving mouse reveals signals from dozens of striatal neurons. **D:** Ca^2+^ signal of an individual neuron (top, green), the average Ca^2+^ signal across all imaged neurons (56 neurons, middle, green) and the simultaneously recorded total body acceleration of a representative mouse (bottom, black). **E:** Color-coded matrix showing activity around movement onset, averaged across all movement onset events (top). Average Ca^2+^ activity across the population of imaged neurons and average total body acceleration across movement onset events are represented by green and black traces, respectively (middle). Peristimulus time histogram (PSTH) of Ca^2+^ events centered around movement onset (bottom). Bin width: 67 ms. **F:** Rates of detected neurons that did (green) or did not (white) significantly modulate their Ca^2+^ signals following movement onset. Shaded areas represent S.E.M.

We began by examining the activity of SPNs and PV interneurons around the onset of self-initiated movement. Six-hundred and thirty seven putative somata were detected in 7 mice [2 imaging sessions per mouse, 45.5 ±33.4 (mean ±SD) neurons per session]. Movement onset and offset times were detected by setting a threshold on the total body acceleration of the mouse [see Methods, Fig. S2 A-B, 171 ±91 (mean ±SD) movements per session]. For each neuron, we calculated the event triggered average (ETA) by averaging the Ca^2+^ signal in a 5 second window around movement onset across all movement events. We found that the average ETA across the population of imaged neurons increased following movement initiation (Fig. 1 D-F). The Ca^2+^ signal around the onset of self-initiated movement followed the total body acceleration and peaked approximately 130 ms after the acceleration peak. The peri-stimulus time histogram (PSTH) centered around the same events of movement onset exhibited a similar peak, indicating that the population of imaged neurons produced more Ca^2+^ events following movement onset (Fig. 1E). Repeating the analysis around movement offset times generated a mirror image of the movement onset responses (data not shown), suggesting that neurons modulate their activity when movement is initiated and that, at the population level, the modulation is maintained throughout the movement. To determine whether the activity of an individual neuron was significantly modulated around movement onset, we devised a bootstrapping-based method (see Methods, Fig. S2 C-E). Twenty seven percent of detected neurons were found to be significantly responsive around the onset of self-initiated movement (Fig. 1F). Thus, while movement responsiveness is evident at the population level, only a subset of the imaged neurons consistently modulate their Ca^2+^ signals around movement onset in a given imaging session.

A closer examination of the ETAs of individual significantly responsive neurons around the onset of self-initiated movement revealed that they exhibit two types of responses (Fig. 1E-F, Fig. S6A-B, G-I): while a majority of responsive neurons (78%) increase their rate of Ca^2+^transients as the total acceleration of the mouse increases (positively modulated, top part of matrix in Fig. 1E & Fig. S3A), some significantly responsive neurons (22%) produce less Ca^2+^transients when the mouse is moving (negatively modulated, bottom part of matrix in Fig. 1E & Fig. S3B). Additionally, movement responses of the imaged neurons were lateralized, displaying a strong preference to contralateral turns over ipsilateral ones (Fig. S7, see Methods). These properties are consistent with known activity patterns of SPNs and PV interneurons during spontaneous movement (Albin et al., 1989; Alexander and Crutcher, 1990; Barbera et al., 2016; Brown, 2007; Chan et al., 2006; Cui et al., 2013; DeLong, 1990; Gerfen et al., 1990; Gittis et al., 2011; Gritton et al., 2019; Hikosaka et al., 2000; Isomura et al., 2013; Jin et al., 2014; JW, 1996; Klaus et al., 2017; Kravitz et al., 2010; Nambu, 2008; Parker et al., 2018; Perk et al., 2015; Tecuapetla et al., 2014).

### Portion of HaR striatal neurons responsive to task-related cues increases systematically as training progresses

To characterize the collective activity of SPNs and PV interneurons during learning, we trained 6 mice in an operant conditioning paradigm to associate an auditory conditioned stimulus (CS) with a reward. Each training session consisted of 42 trials. In each trial, an auditory cue (CS+ or CS–) was presented for 10 seconds in a pseudo-random manner. To receive a sucrose-water reward, the freely moving mouse was required to enter its head into the reward delivery port while the CS+ was being presented. A head-entry during CS– presentation was not rewarded (Fig. 2A).

**Figure 2.**
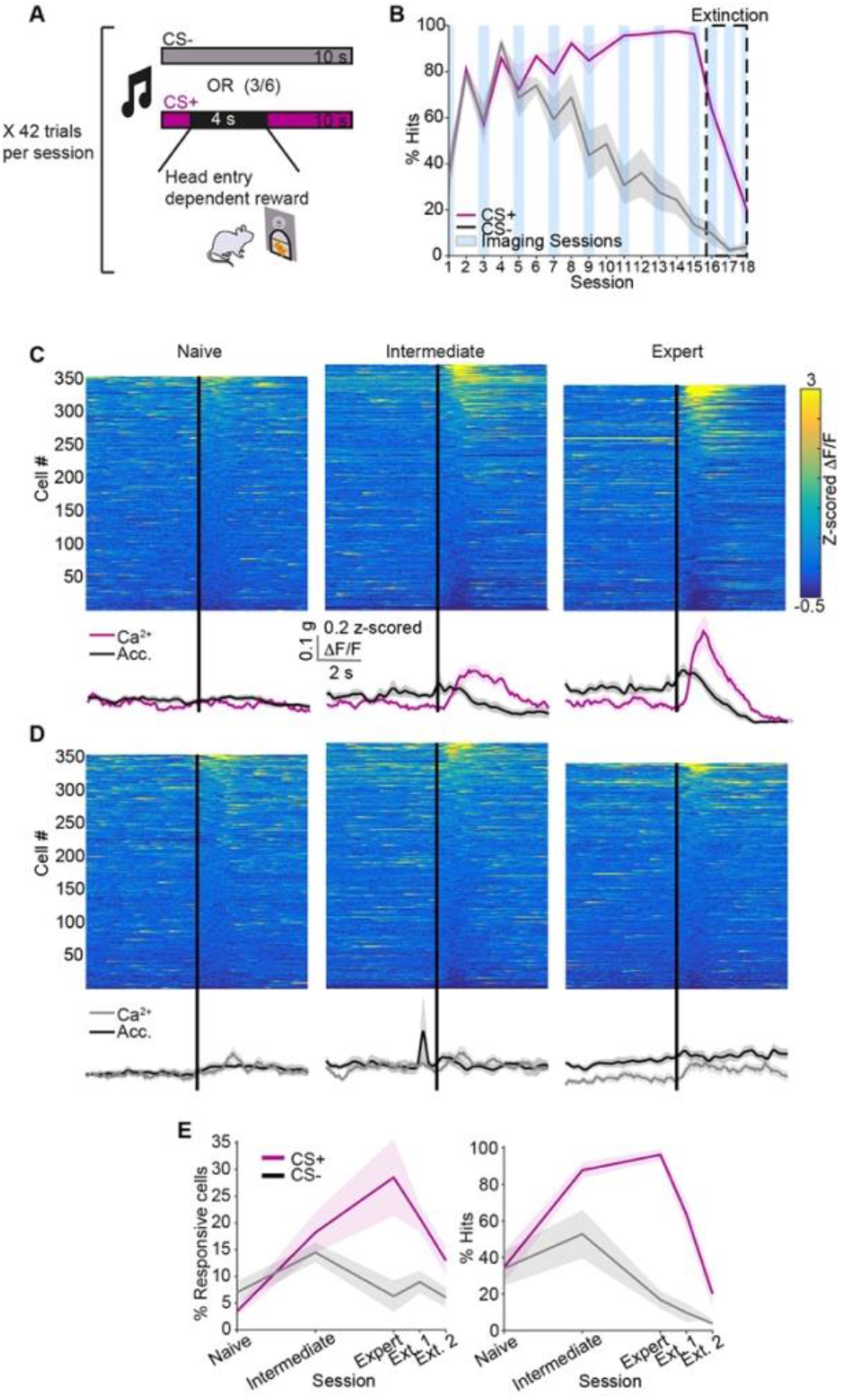
Responses to task-related cues develop with training. **A:** Mice were trained in an operant conditioning paradigm to associate an auditory CS with a sucrose-water reward. Each session consisted of 42 trails. In each trial, a cue (CS+ or CS–) was presented for 10 seconds in a pseudo-random manner. If the mouse entered its head into a designated port during the 10 second window following CS+ (but not CS–) onset, reward was available for 4 seconds. **B:** The learning rate quantified as the portion of cue presentations during which the mouse entered its head into the port (% hits). Blue rectangles mark imaging sessions and the dashed box marks extinction sessions. **C:** Color-coded matrix showing neuronal activity around CS+ presentation averaged across all CS+ presentation events for naïve (left), intermediate level (middle) and expert (right) mice. Average Ca^2+^ activity across the population of imaged neurons and average total body acceleration across CS+ presentation events are represented by pink and black traces, respectively (bottom).**D:** Same as C, for CS– presentation. Average Ca^2+^ activity across the population is represented by gray traces. **E:** Rate of responsive neurons (left) and ‘hit’ rate (right) around CS+ (pink) and CS– (gray) presentation on various stages of training. Shaded areas represent the S.E.M.

Mice successfully acquired the association between the CS+ and the sucrose-water reward (Fig. 2B). To quantify learning, we measured for each cue (CS+ and CS–) the portion of cue presentations during which the mouse entered its head into the port attempting to obtain the reward (‘hits’). On advanced training days, mice perform head-entries upon almost every CS+ presentation, while the rate of CS– ‘hits’ decreases to ~15%, indicating that the CS+ was associated with the sucrose-water reward while the CS– was not. Once the association was established, mice underwent three extinction sessions. Those sessions were identical to conditioning sessions, but no reward was delivered. The association between the CS+ and the reward was quickly extinguished and mice ceased to perform head-entries.

To characterize the collective activity in various stages of learning, we compared ETAs in a 10 second window around cue presentation in naïve, intermediate level and expert mice (see Methods). We considered one session per mouse at each learning stage and detected a total of 355, 342 and 373 putative somata on naïve, intermediate and expert training sessions, respectively. In naïve mice, the average Ca^2+^response following CS+ presentation was weak. The population response developed as training progresses and was strong on advanced conditioning sessions when mice were experts at the task (Fig. 2C). In contrast, average CS– responses remained weak throughout training (Fig. 2D). This was evident in the average ETAs representing the population signal, as well as in the ETAs of individual neurons. Importantly, CS+ responses were not determined by the total body acceleration of the mouse or by reward delivery (Fig. S4).

The event rates of neurons significantly responsive to CS+ or CS–tightly corresponded to task performance (Fig. 2E). In naïve mice, very few neurons responded around either cue. In intermediate level mice, towards the middle of training, there was an increase in the portion of significantly responsive neurons for both CS+ and CS– presentations. On advanced conditioning sessions, when mice were experts at the task, a larger portion of detected neurons significantly modulated their activity following CS+ presentation, while very few responded to CS– presentation. This profile echoed the progression of learning as represented by the rates of CS+ and CS– ‘hits’, suggesting that the activity of these neurons is meaningful in the context of the task. Notably, while both the average population signal and rate of significantly responsive neurons increase systematically as training progresses, even in expert mice, only a subset (~30%) of detected neurons significantly modulated their activity in response to CS+ presentation.

Similarly to movement-responsive neurons, neurons that significantly modulated their signals around task-related cues could be divided into two functional groups – positively and negatively modulated neurons (Fig. 2 C-D & Fig. S5). Here too, the majority of responsive neurons increased the rate of their Ca^2+^ transients following cue presentation, while the rest exhibited less Ca^2+^ events. Thus, subsets of SPNs and PV interneurons modulate their activity around self-initiated movements as well as task-related cues in an operant conditioning paradigm.

### Individual HaR striatal neurons are recruited to respond to the same behavioral events in a non-reliable fashion across and within imaging sessions

We next examined the reliability of neuronal responses around various behavioral events. Will a neuron that is significantly responsive around a specific behavior in a given imaging session continue to code for it on subsequent sessions? Our experimental approach allowed us to track individual neurons across multiple imaging sessions. To examine the stability of movement responses, we detected neurons that were active on two free movement sessions. Out of 88 such neurons, 22% significantly responded around movement onset on the first session, 27% responded on the second session, and only 9% significantly modulated their signals around self-initiated movement on both free movement sessions (Fig. 3C). While the 9% overlap is greater than the expected rate if neurons are randomly recruited to respond around movement onset (e.g., 6%), the identity of significantly responsive neurons appears to be highly dynamic across imaging sessions.

**Figure 3.**
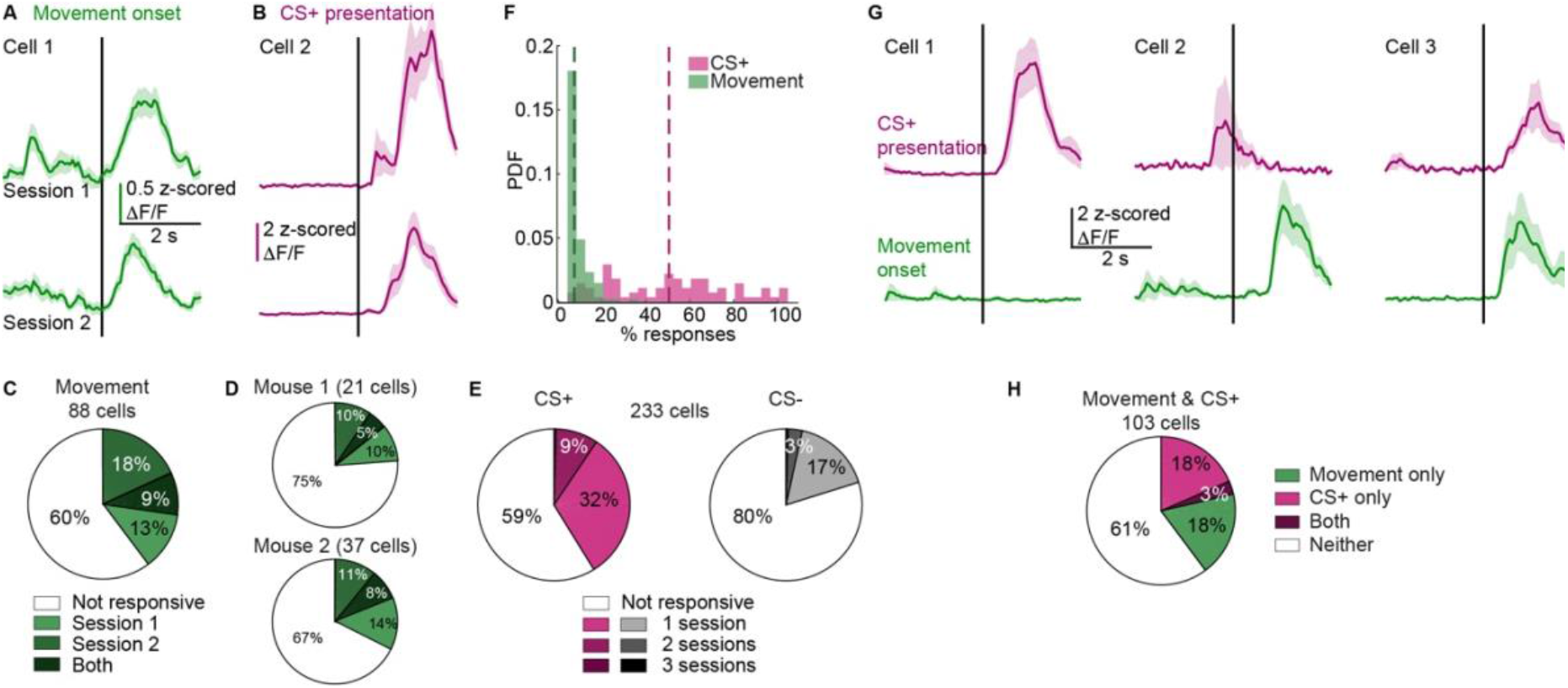
Responses around movement onset and cue presentation are dynamic across imaging sessions. **A:** Average Ca^2+^ signal of an example neuron on the first (top) and second (bottom) free movement imaging sessions that exhibited significant responses around movement onset on both sessions. **B:** Same as A, around CS+ presentation. **C**: Percentage of neurons significantly responsive on the first (light green) or second (medium green) free movement imaging sessions, both of them (dark green) or neither (white). **D:** Percentage of neurons from mouse 1 (top) and mouse 2 (bottom) significantly responsive on the first (light green) or second (medium green) free movement imaging sessions, both of them (dark green) or neither (white). **E:** Percent of neurons significantly responsive on one (light), two (medium) or three (dark) advanced conditioning sessions around CS+ (left, pink) and CS– (right, gray) presentation. **F:** PDF of the percent of movement onset (green) and CS+ presentation (pink) events after which the neuron produced a Ca^2+^ event for positively modulated neurons. **G:** Examples of neurons exhibiting significant and non-significant responses (according to our bootstrapping criterion, see Methods) around movement onset and CS+ presentation. Cell 1 responded significantly only to CS+ presentation. Cell 2 responded significantly only to movement onset. Cell 3 responded significantly to both. **H:** Percentage of neurons significantly responsive around movement onset (green), CS+ presentation (pink), both (dark purple) or neither (white) out of the neurons that were detected both in a free movement session and in an advanced conditioning session.

In 5 of the 7 imaged mice, the two free movement sessions were several weeks apart. To test if this large interim could account for the high variability in the populations of responsive neurons, we focused on the two mice whose free movement sessions were only a few days apart (Fig. 3D). This analysis revealed very similar percent overlap between the populations of neurons significantly responsive on the two sessions, indicating that the low reliability is not due to changes occurring slowly over time.

To assess the reliability of the responses around task-related cues, we examined the signals of neurons that were detected on three advanced training sessions (2-4 days apart) around the presentation of the CS+ and CS– (Fig. 3E). Of 233 detected neurons, 32% significantly responded to CS+ presentation on a single session, 9% were responsive on two sessions, and only one neuron significantly modulated its signal following CS+ presentation on all three sessions. Similarly, 17% of the neurons were significantly responsive around CS– presentation on a single session, 3% were responsive on two sessions, and one neuron was responsive on all three sessions. These percentages of overlap are consistent with neurons being randomly and independently recruited to respond around cue presentation on different imaging sessions.

An important caveat is that the movement events that we considered, detected by setting a threshold on the total body acceleration, can include a wide range of movements. Thus, the changes in the identity of significantly responsive neurons between the first and second free movement sessions could be due to differences in the behavior of the mice. That being said, while still varied, the behavior around task-related cues, especially on advanced training days, is much more consistent in terms of the acceleration profile (Fig. S6). Our analysis of CS+ and CS– responses across imaging sessions indicates that while the percentage of significantly responsive neurons increases with training, their identity is dynamic and shifts from session to session.

To examine the reliability of the responses within a given session, we considered significantly responsive neurons whose signals were positively modulated around movement onset (Fig. 3F). Positively modulated neurons are the majority of significantly responsive neurons and tend to generate more Ca^2+^ events following movement onset compared to prior to it (Fig. S3). For every such neuron, we calculated the portion of movement initiations after which it produced a Ca^2+^ event. Interestingly, even neurons that were categorized as positively modulated for a given imaging session only produced Ca^2+^ events in an average of 7.2% of movements. Repeating the analysis for neurons that were positively modulated around CS+ presentation (Fig. 2D & Fig. S5) produced a higher percent overlap, with neurons responding with a Ca^2+^ event for an average of 46.4% of CS+ presentations. The higher reliability of the responses following CS+ presentation compared to movement onset may reflect a difference in the organization of striatal neurons around spontaneous movement and learned behaviors. Alternatively, the disparity may be due to the lower number of cue presentation events (23 cue presentations per session compared to an average of 171 movement initiation events), as well as the more homogeneous behavior associated with these events (Fig. S6). Nevertheless, our results suggest a low reliability in the responses of SPNs and PV interneurons around both types of behavioral events.

### Individual HaR neurons are recruited to respond to diverse actions in an independent fashion

Imaged neurons significantly modulate their activity following the onset of self-initiated movement and task-related cues (Fig. 1 & Fig. 2). Do the same neurons respond to the different behavioral events, or are there distinct subpopulations designated to code for each behavior? To address this question, we tracked individual neurons across multiple imaging sessions and examined their responsiveness around movement initiation and CS+ presentation.

One hundred and three neurons were detected on both a free movement session and an advanced conditioning session. Of these neurons, 21% exhibited significant responses around movement onset only, 21% significantly modulated their activity following CS+ presentation only, and 3% were responsive around both movement initiation and cue presentation (Fig. 3H). The percent of neurons significantly modulating their signals around both behavioral events is consistent with neurons being randomly and independently recruited to respond to movement initiation and CS+ presentation.

Taken together, our results indicate that while SPNs and PV interneurons consistently modulate their activity around self-initiated movement and CS+ presentation on the population level, the identity of responsive neurons is not maintained. Rather, neurons are dynamically recruited to code for these different behavioral events on separate occurrences.

### Signal correlations between pairs of HaR neurons are independent of distance

Are responsive neurons selected at random with each repetition of the behavior? Or is there a more complex internal organization that our analysis is unable to capture, perhaps due to the vague definition of the behavior we are examining (e.g., movement initiations)? Recent studies have shown that SPNs form spatially compact neural clusters in which nearby neurons are co-active and code for the same type of movement (Barbera et al., 2016; Parker et al., 2018; Shin et al., 2020). Notably, this influential conclusion was drawn from the observation that the correlations between pairs of SPNs decay with distance. Thus, to test whether the unreliability of individual responses in striatal HaR neurons is compensated for by co-activation, we calculated pairwise signal correlations among our imaged neurons and the dependence of signal correlations on the distance between the neurons’ centers.

To test whether nearby neurons tend to exhibit similar activity patterns, we calculated signal correlations between the Ca^2+^ traces of pairs of neurons during rest and movement. When all simultaneously imaged neuronal pairs were included, the average correlation (averaged over 10 μm distance bins and then across bins, see Methods) is very low during both rest and movement (0.004-0.005, Fig. 4A, C, Fig. S7C). When positively and negatively modulated neurons are considered separately, pairwise correlations during both movement states are larger by an order of magnitude and slightly (but not significantly) larger during movement compared to during rest (Fig. 4B,D). Notably, all of these average values are considerably lower than what was previously reported (Barbera et al., 2016; Shin et al., 2020) (see Discussion). Strikingly, for all functional groups and both movement states, pairwise correlations did not depend on the distance between the neurons’ centers (Fig. 4A-D).

**Figure 4.**
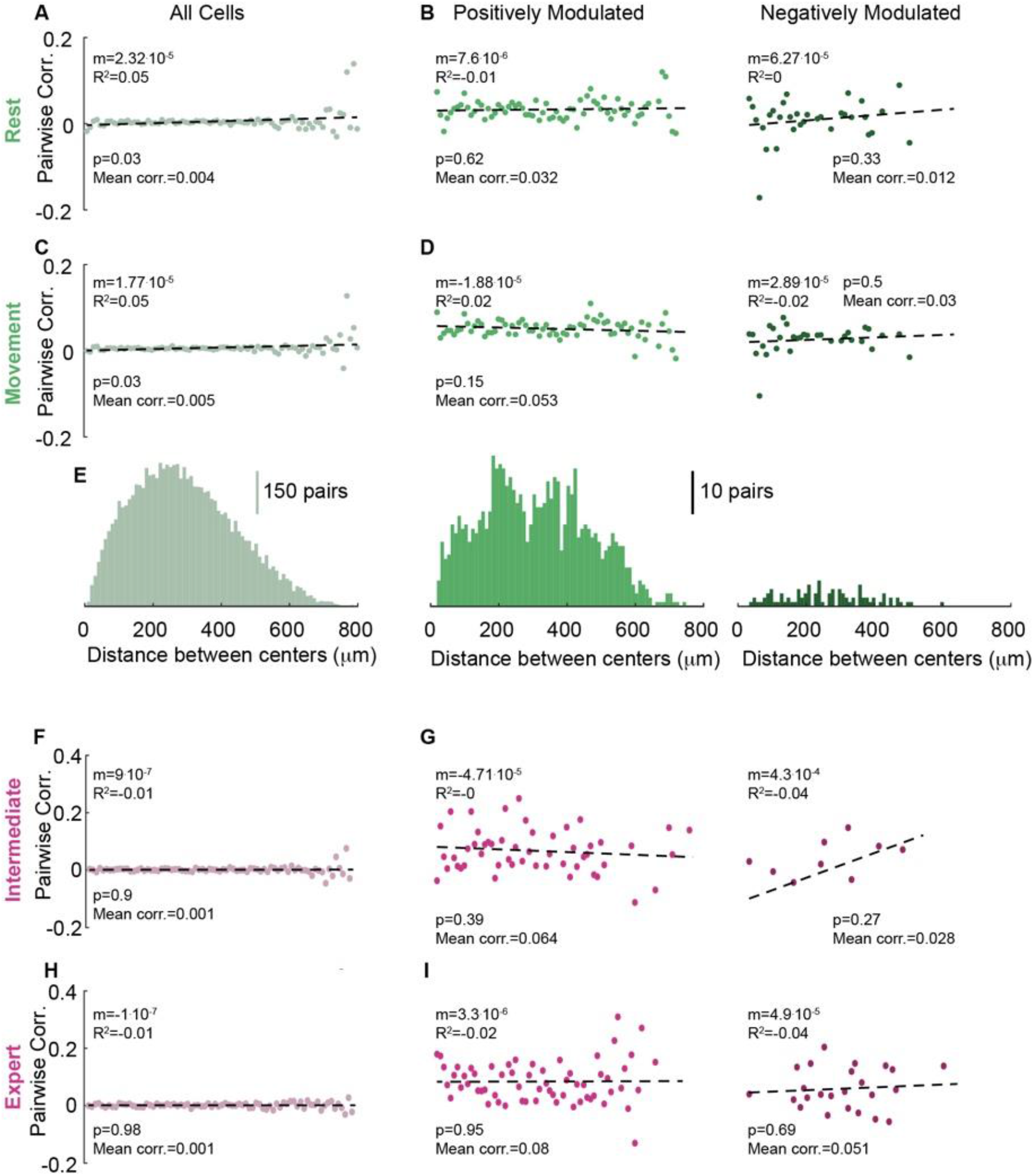
Pairwise correlations are not distance dependent. **A:** Pairwise correlations as a function of the distance between the neurons’ centers for all pairs of co-imaged neurons during rest. Each point is the average correlation across all neuronal pairs belonging to the relevant distance bin. Dashed line is the linear fit to the data points. **B:** Same as A, divided into positively (left) and negatively (right) modulated pairs. **C:** Same as A, around movement onset. **D:** Same as B, during movement. **E:** Distribution of neuronal pairs for the various distances for all pairs (left), positively (middle) and negatively modulated pairs (right). Bin width is 10 μm. **F:** Same as B, around CS+ presentation on intermediate training session. **G:** Same as F, divided into positively (left) and negatively (right) modulated pairs. **H:** Same as F, on expert training session. **I:** Same as G, on expert training session.

Next, we calculated the signal correlations between the Ca^2+^ traces of neuronal pairs around CS+ presentation. Similarly to movement responses, when all pairs were included, pairwise correlations around cue presentation were very low on both intermediate and expert training sessions (0.001, Fig. 4F, H). When positively and negatively modulated neurons were considered separately, pairwise correlations were increased by an order of magnitude. Importantly, for CS+ responses, at both stages of training and for all functional groups, pairwise correlations did not significantly depend on the distance between the neurons, either. To conclude, we observe weak signal correlations between pairs of our imaged neurons, challenging the idea of stable, consistently co-activated cell clusters. These correlations are also distance independent for both self-initiated movement and CS+ presentation, in contrast to previous findings of spatially compact SPN clusters (Barbera et al., 2016; Shin et al., 2020). Interestingly, correlations in the background neuropil activity, represented by the Ca^2+^ signals in the annuli surrounding the various somata (see Methods), did exhibit a clear distance dependence with pairwise correlations decreasing sharply between 0 and 200 μm (Fig. S7B, see Discussion).

### Trial-by-trial covariation among HaR neurons around movement onset are attributable to variability in behavior

Signal correlations between neuronal pairs reflect the degree to which their average responses are modulated together by the sensory or behavioral event. However, signal correlations do not capture the role of the ongoing moment-by-moment synergy between individual neurons that may underlie dynamic cell assembly formation. To investigate the propensity of the imaged neurons to be co-active beyond common increases or decreases in the average rate of Ca^2+^transients, we calculated joint peristimulus time histograms (JPSTHs) for the various neuronal pairs (Aertsen et al., 1989; Gerstein and Perkel, 1969; Joshua et al., 2009). The JPSTH is a measure of noise correlations or the degree to which trial-by-trial fluctuations in the response are shared by a pair of neurons. It is derived by subtracting the shift predictor matrix from the product of the PSTHs of the two neurons (Aertsen et al., 1989) (see Methods), resulting in an estimate of unpredicted correlations.

To assess noise correlations in the responses to self-initiated movement, we calculated the JPSTHs for pairs of neurons around movement onset. The JPSTHs of all neuronal pairs were then normalized and averaged to derive the population JPSTH (Aertsen et al., 1989; Joshua et al., 2009) (see Methods). The diagonal of the resulting matrix quantifies the average time varying modulation of zero-lag noise correlation across the population (Fig. 5A). Noise correlation values were similar along the entire diagonal, except for the 1-second window immediately before movement initiation showing a correlation drop (Fig. 5A). This reduction is likely due to the decreased acceleration and neuronal activity that mark the period leading up to movement onset (Fig. 1E).

**Figure 5.**
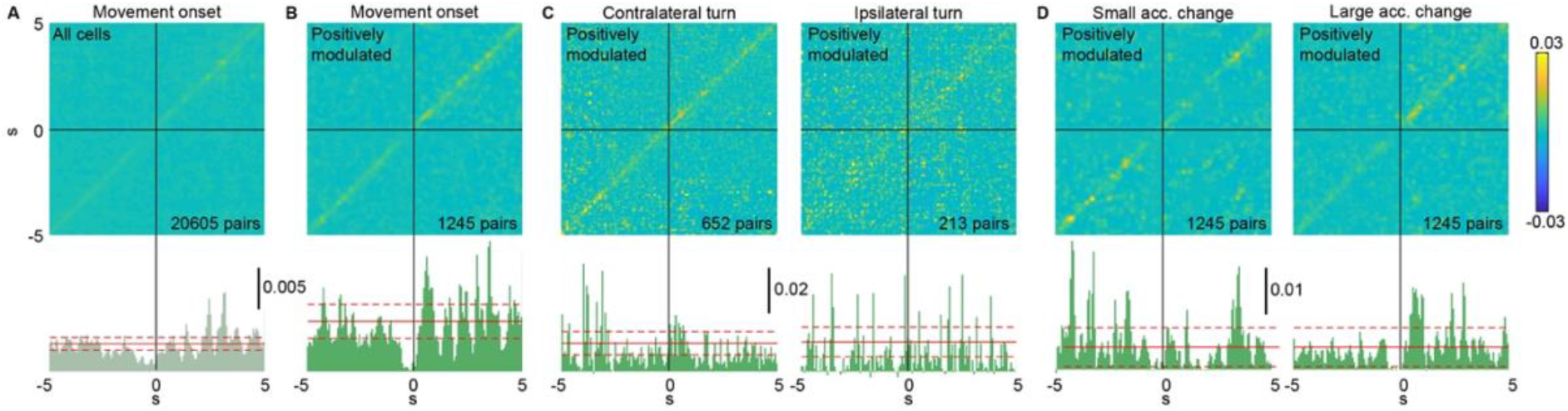
Movement-related trial-by-trial correlations are strongly related to behavior variability. **A:** Population joint peristimulus time histogram (JPSTH), averaged across all neuronal pairs, centered around movement onset (top). Bar plot shows values of the JPSTH diagonal (bottom). Solid red line represents the mean value of the JPSTH diagonal in a 10 second time window far removed from movement onset events. Dashed red lines represent the mean ±the SD. **B:** Same as A, for pairs of positively modulated neurons. **C:** Same as B, for contra-(left) and ipsilateral (right) turns. **D:** Same as B, for movement onset events with a small (left) or large (right) acceleration change.

Next, the analysis was repeated, considering only pairs of positively modulated neurons. Focusing on this population resulted in a larger peak in noise correlation immediately following movement initiation (Fig. 5B). The diagonals of the population JPSTHs for pairs of negatively modulated neurons or pairs with opposite modulation directions did not exhibit significant peaks (data not shown).

Movement onset times were detected by setting a threshold on the total body acceleration of the mouse and therefore include a wide range of movements. To test whether joint trial-by-trial variations in neuronal activity arise from variability in the mouse’s behavior, we repeated the analysis for subsets of more stereotyped movement events. Limiting the analysis to contra- or ipsilateral turns eliminated the peak in the diagonal of the population JPSTH (Fig. 5C). Finally, the same calculations were conducted focusing separately on movements with small and large acceleration changes (Fig. 5D). While the JPSTH diagonal for movements accompanied by a small acceleration change exhibits a small peak, which does not exceed random fluctuations in the values of the diagonal, movements with large acceleration changes induce a stronger peak in noise correlation. Taken together, these results suggest that trial-by-trial correlations around the onset of self-initiated movements are mostly due to variability in the movement of the mouse driving a temporal covariation among neurons in the assembly across repeated events.

### Trial-by-trial correlations following the presentation of task-related cues

To investigate trial-by-trial correlations during learning, we calculated the JPSTHs of neuronal pairs around CS+ presentation focusing on the expert training session and the first extinction session. When all neuronal pairs were included, the diagonals of the population JPSTHs exhibited no peaks for any stage of training (data not shown). Similarly to the movement onset JPSTH, focusing on pairs of positively modulated neurons resulted in significant noise correlation peaks following CS+ presentation in expert mice (Fig. 6A). Importantly, mouse behavior following CS+ presentation, particularly at advanced stages of training, is relatively uniform. First, mice successfully obtain the sugar-water reward in the vast majority of trials during the expert training session (128 out of 137 trials in 6 trained mice). Additionally, mice acceleration profiles following CS+ presentation become less variable as training progresses (Fig. S6). Next, the analysis was repeated focusing on the first 10 extinction trials. These trials were unrewarded, like the rest of the extinction session, but they occurred early enough in the session so that the association between the cue and the reward had not yet been lost. In this case as well, noise correlations rise shortly after the presentation of the CS+ (Fig. 6B). To further reduce the variability in the behavior, we divided expert session trials into those accompanied by small (1^st^ quadrant) and large (4^th^ quadrant) acceleration changes (Fig. 6 C-D). In both cases, the diagonals of the population JPSTHs maintain their peaks following CS+ presentation.

**Figure 6.**
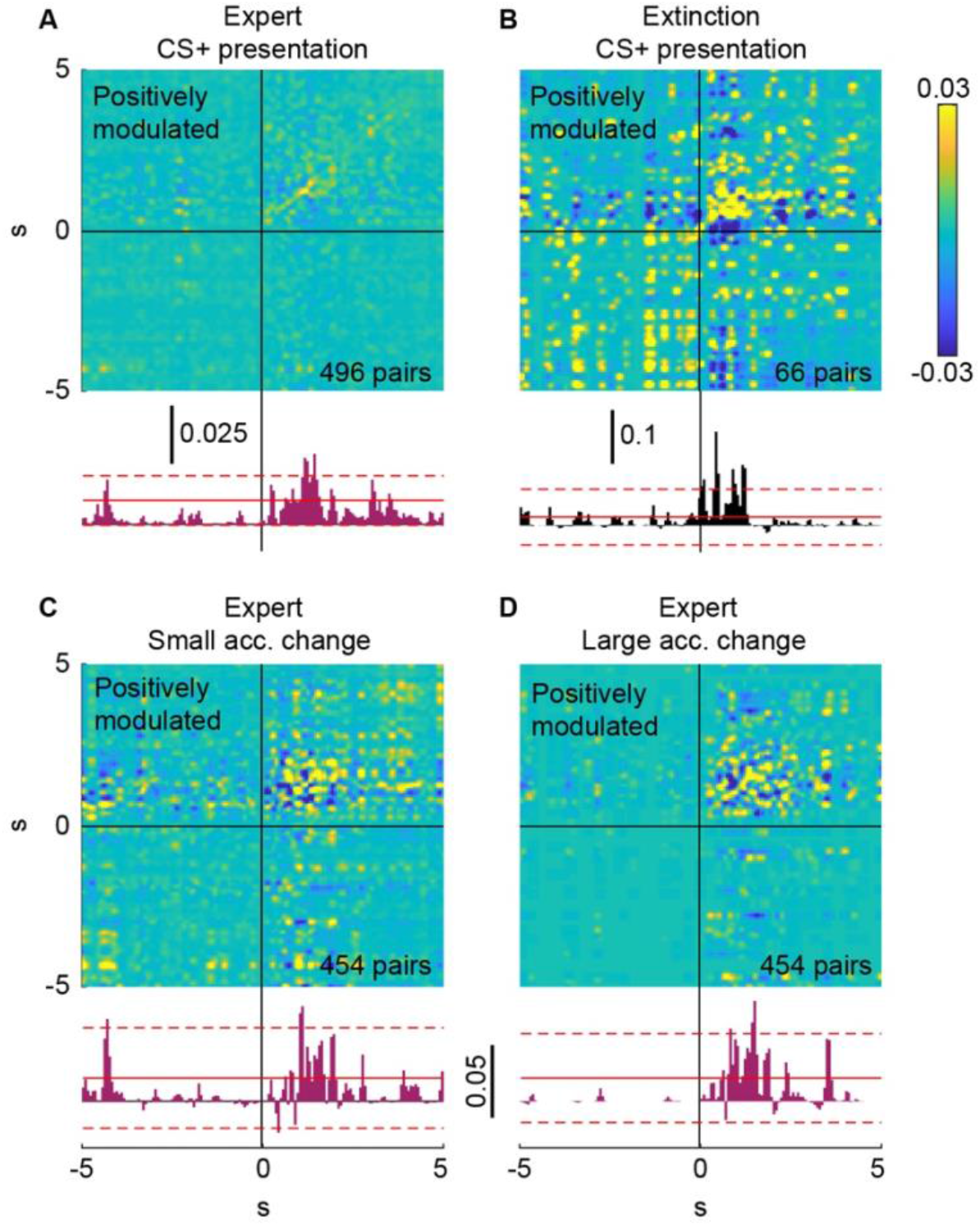
Trial-by-trial correlations following CS+ presentation. **A:** Population JPSTH, averaged across pairs of positively modulated neurons, centered around CS+ presentation on expert training session (top). Bar plot shows values of the JPSTH diagonal (bottom). Solid red line represents the mean value of the JPSTH diagonal in a 10 second time window far removed from movement onset events. Dashed red lines represent the mean ±SD. **B:** Same as C, for first 10 trials of the first extinction session. **C:** Same as A, for CS+ presentation events accompanied by a small change in acceleration. **D:** Same as A, for CS+ presentation events accompanied by a large change in acceleration.

In summary, while trial-by-trial correlations around the onset of self-initiated movements seem to arise from variability in the behavior of the mouse, noise correlations surrounding the presentation of task-related cues persisted when we examined subsets of behavioral events with more similar characteristics. Thus, while evidence for dynamic formation of correlated assemblies on individual actions is weak, if it occurs it seems to occur for learned behaviors (of acquired value) but not for innate self-initiated behaviors.

## Discussion

In this study, we aimed to define the response reliability of HaR striatal neurons in freely moving animals during both self-initiated and learned actions. To this end, we conducted microendoscopic Ca^2+^ imaging in freely moving mice during spontaneous movement and at different stages of learning in an operant conditioning task. The Ca^2+^ indicator GCaMP6f was expressed in a sparse neuronal population consisting predominantly of SPNs, with PV interneurons making up a smaller but substantial portion (Fig. S1). These subtypes represent the HaR striatal populations that receive a small number of strong afferents from the cortex (Plotkin et al., 2011) and thalamus (Johansson and Silberberg, 2020). Because the sum of fewer inputs is expected to exhibit more variability in spike timing, we predicted and found that HaR neurons would exhibit low levels of response reliability.

At the population level, imaged neurons showed clear responses to both self-initiated movement and the presentation of task-related cues. However, an examination of the signals of individual neurons revealed that a relatively small subset of neurons (approximately 30%) significantly modulated their activity following these behavioral events, while in most neurons, the rate of Ca^2+^ transients was not significantly altered (Fig. 1F, Fig. 2E). This responsiveness rate is consistent with the findings in a recent electrophysiological study, where a similar proportion of SPNs modulated their signals near starts and ends of locomotion in freely moving mice (Fobbs et al., 2020). Previous studies quantifying the portion of SPNs coding for changes in value or for cue presentation in conditioning tasks also reported comparable percentages (between 25 and 50%) (Bakhurin et al., 2016; Shin et al., 2018). Thus, during both spontaneous movement and learning, the striatal HaR response depends on a small subset of the neurons – a functional organization that is likely to give rise to highly variable processing, in accordance with our observations (Fig. 4).

Our mice expressed GCaMP6f in a sparse population of striatal neurons (Fig. S1). This sparse expression enabled us to track and compare the responses of the same individual neurons across sessions, days, and even weeks apart. While the proportion of movement responsive neurons remained relatively stable across free movement sessions, a large fraction of neurons that significantly modulated their activity around movement initiation on the first session did not do so on the second session (Fig. 3A,C,D). Similarly, the rate of neurons that significantly responded to CS+ presentation gradually increased as training progressed and was stable on advanced training sessions (Fig. 2). However, the identity of significantly responsive neurons varied considerably from session to session (Fig. 3B, E). These findings, which were facilitated by the sparse expression of the indicator, suggest that the recruitment of striatal neurons to respond to various behavioral events is dynamic. In addition, the identity of neurons significantly modulating their activity around movement initiation, cue presentation or both of them is consistent with neurons being randomly recruited to respond to various behavioral events (Fig. 3H). Taken together, our results suggest that action representation in HaR striatal neurons, which receive strong cortical and thalamic inputs, changes dynamically within and among sessions through the moment-by-moment formation of distinct cell assemblies.

What are the advantages of dynamically encoding a population response reliably while a majority of neurons do not participate in the encoding and while the identity of the neurons encoding constantly changes? One possibility is that the question is ill posed because we are not tracking enough parameters of the outside world. By collapsing all the neurons onto a single coarse behavior (e.g., movement initiations) we might be missing other dimensions being encoded (Liberti et al., 2022). While we cannot rule out that possibility, it is also possible that there are inherent advantages in dynamic encoding. The striatum is widely viewed as essential for selecting the appropriate motor plan during ongoing behavior. To accomplish this task in a constantly changing environment, it is critical for the responses of striatal neurons, and particularly those of projection neurons, to be highly flexible. A scheme in which the identity of responsive neurons changes from day to day and even between trials within the same day could simplify the encoding of new stimulus-action contingencies.

Alternatively, our data are consistent with the striatum implementing a population code, where the portion of the population activated around a behavioral event is stable, and the identities of the responsive neurons change continuously. Because each pallidal and/or nigral neuron receives afferents from many SPNs (Percheron et al., 1984), it may be indifferent to the identity of the projecting neurons. Similarly, while PV interneurons are few in the striatum, their dense axonal arborization extends far from the soma, with each PV interneuron synapsing onto hundreds of SPNs (Berke, 2011; Koos et al., 2004; Tepper, 2010). Thus, while it may be that the low response reliability of striatal neurons does not improve the information decoding in downstream structures, it also does not degrade it due to the highly convergent architecture of these neurons’ BG circuits (Percheron et al., 1987).

Next, theorizing that the unreliable responses might be compensated for by co-activation, we exploited our ability to simultaneously image the activity of dozens of striatal neurons and explored their signal and noise correlations in different behavioral states. We found that pairwise signal correlations in HaR striatal neurons are independent of the distance between them, with similar results during movement and rest, as well as near cue presentation at different stages of conditioning (Fig. 4). These findings are in line with previous *in vitro* studies, which demonstrated that nearby SPNs (making up the majority of our neurons) receive very different inputs (Kincaid et al., 1998; Zheng and Wilson, 2002) and therefore should not be more correlated than distant ones.

While consistent with striatal connectivity, the distance independence of pairwise correlations (Fig. 4) in our mixed population, which comprises a majority of SPNs (Fig. S1), contrasts with prior reports of spatially compact SPN clusters in the striatum (Barbera et al., 2016; Parker et al., 2018; Shin et al., 2020). Importantly, these previous results were derived through the simultaneous microendoscopic imaging of thousands of SPNs, likely also including a substantial background neuropil signal. Neuropil signals in other striatal subtypes have been found to exhibit high spatial correlations and show different kinetics from somatic signals (Legaria et al., 2021; Rehani et al., 2019) – a result that we reproduced here for HaR striatal neurons (Fig. S7). Additionally, background neuropil correlations in our data show a strong distance dependence, decaying steeply as the distance grows from 0 to 200 μm (Fig. S7B). Source separation is challenging in a dense neuronal population. It is thus possible that the discrepancy between our observations and previous results could emerge from spatially correlated neuropil activation contaminating somatic signals in a dense SPN population and introducing spurious, distance dependent correlations.

Further supporting our proposed explanation, pairwise signal correlations in striatal HaR neurons were, on average, weaker by orders of magnitude than previously reported values (Fig. 4). Because these studies used both slow and fast GCaMP6 variants (Barbera et al., 2016; Shin et al., 2020), it is unlikely that the discrepancy stems from differences in the kinetics of the indicator. However, if previously reported somatic correlations include spurious correlations originating from the neuropil signal as we suggest, and if the sparse expression generates a weaker background neuropil signal, then we expect the correlation values in our data to be lower. We propose that the sparse expression of the Ca^2+^ indicator in our mice (Fig. S1) produces a weak neuropil signal compared to those arising from densely expressing populations. As a result, the background neuropil signal introduces less spurious correlations between somata, decreasing pairwise correlation values and eliminating their distance dependence. While we cannot fully address this point based on our data, which do not target SPNs exclusively, we believe that the concerns raised by our results merit consideration.

Finally, we investigated trial-to-trial correlations between pairs of striatal HaR neurons by calculating their JPSTHs around spontaneous movement onset and cue presentation. For self-initiated movements, we found that any covariation beyond common modulations in the average rate of Ca^2+^ transients is most likely explained by variability in the movement itself, rather than the dynamic organization of responsive neurons into a cluster around that movement (Fig. 5). However, with respect to learned associations, the JPSTH analysis suggests that there may be additional components of a dynamic organization into co-active assemblies (Fig. 6).

In conclusion, our analysis of sparsely-labeled SPNs and PV interneurons, which represent striatal HaR neurons, in freely moving mice demonstrated that individual neurons are recruited in a stochastic and dynamic fashion to respond to specific actions, presenting low response reliability. The JPSTH analysis suggests inherent differences in the collective dynamics during learning and innate behaviors, possibly signifying that the coactivation during task learning is more “meaningful” to the neural code than around spontaneous movement. We speculate that the low degree of response reliability affords the SPNs more flexibility in responding to a battery of potential actions and/or various multiplexed encodings. On the other hand, because of the large degree of convergence in the pallidum and substantia nigra pars reticulata (Bolam et al., 1993; Percheron et al., 1987, 1994), this lack of reliability will not affect the constancy of downstream pallidal or nigral representations.

## Methods

### Animals

Experimental procedures adhered to and received prior written approval from the Hebrew University’s Institutional Animal Care and Use Committee (protocol number MD-19-15856-4). Two-to-seven-months-old mice were used in the experiments. Animals were housed in a reversed light-dark facility.

Transgenic mice that express Cre-recombinase sparsely in the striatum (stock number 006410; Jackson Laboratories, Bar Harbor, ME, USA) were cross-bred with Cre-dependent, Tet-controllable, GcaMP6f (Ai148, TIT2L-GC6f-ICL-tTA2) mice (stock number 030328; Jackson Laboratories, Bar Harbor, ME, USA) to generate a sparse GCaMP mouse. The population of GcaMP6f-expressing neurons was dominated by SPNs (~60%), with a moderate GABAergic interneuron population (~30%) consisting mostly of PV interneurons, and a smaller proportion of CINs (Fig. S1).

### Gradient refractive index lens implantation

Eight-to-fourteen-week-old mice were deeply anesthetized with isoflurane in a non-rebreathing system and placed in the stereotaxic frame. Temperature was maintained at 35°C with a heating pad, artificial tears were applied to prevent corneal drying, and animals were hydrated with a bolus of injectable saline (5 ml/kg) mixed with an analgesic (5 mg/kg carpofen). A bolus of 33 mg/kg of a ketamine-xylasine mixture was injected initially to stabilize the preparation for induction of anesthesia. A hole slightly wider than the 1 mm diameter (4 mm long) gradient refractive index (GRIN) lens was drilled into the skull in aseptic conditions. The hole was centered around the following coordinates: anteroposterior, +0.5 mm; mediolateral, +2.3 mm; and dorsoventral, −2.8 mm, relative to bregma using a flat skull position. We aspirated cortex with a 27-30 G needle to a depth of approximately 2 mm (just past the corpus callosum) and then fit the lens in snugly. We next applied C&B-Metabond (Parkell, Edgewood, NY, USA) to cement the lens into place together with a head bar for restraining the mouse when necessary. After lens implantation, mice were housed individually under a reversed light-dark cycle. Two to three weeks later we attached a baseplate to guarantee that the microendoscope will be properly aligned with the implanted GRIN lens. To ensure opacity to light, the Metabond was painted with black nail polish.

### Microendoscopic imaging

Microendoscopes (nVista, Inscopix, Palo Alto, CA, USA) sampled an area of approximately 600 × 900 μm (pixel dimension: 1.2 μm) at 15 frames/s. Movies were motion corrected with the Inscopix Data Processing Software suite (IDPS, Inscopix, Palo Alto, CA, USA). The constrained non-negative matrix factorization (CNMF-E) algorithm (Pnevmatikakis et al., 2016; Zhou et al., 2018) was applied to motion corrected movies to detect putative somata and extract traces of fluorescence changes over time. False-positives in the CNMF-E output (i.e. non-neuronal spatial footprints and fluorescence traces) were filtered out of the final dataset manually. Data from one mice with weak transfection or from sessions from all mice with too few somata were discarded. The identification of the same neurons across multiple imaging sessions was performed using the probabilistic approach for cell registration (Sheintuch et al., 2017).

For each somatic region of interest (ROI), detected using CNMF-E, an annulus was defined as a ring of pixels with the same area as the somatic ROI, whose inner diameter is the distance of the point on the border of the ROI that is farthest from its center-of-mass plus 5 additional pixels. In annular ROIs, we extracted fluorescence changes over time (ΔF/F) such that *ΔF/F*= 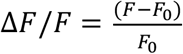
,where F represents the raw fluorescence recorded and F0 denotes the minimal averaged fluorescence across 2 second periods (with 1 second long overlaps) throughout the measurement.

### Behavioral paradigm

In each imaging session, mice were head restrained on a running saucer, mounted with a microendoscope and placed in a behavior chamber (21 cm *×* 18 cm *×* 13 cm) lit by diffuse infrared light, where they could move freely. Naïve mice underwent two free-movement imaging sessions, in which their spontaneous movements were monitored. Next, mice were placed under a food restriction regime, and their weight was maintained at approximately 85% of its initial value throughout training. For two hours before habituation or training sessions mice were also water-deprived. Each mouse underwent three habituation sessions, in which it was required to enter its head into a designated port in the behavior chamber and consume sucrose-water (30% sucrose). Head entries were detected using an infrared photobeam sensor (Med associates, Fairfax, VT, USA). The session ended after 30 minutes, or once the mouse successfully consumed the sucrose-water 10 times.

Habituated mice underwent an operant conditioning paradigm in which an auditory conditioned stimulus (CS) was associated with a sucrose-water reward. Each conditioning session consisted of 42 randomly spaced trials (an average of 30 or 60 seconds between consecutive trials). In each trial, one of two auditory cues (CS+ or CS–; 3200 Hz and 4300 Hz, respectively) was presented in a pseudo-random manner for 10 seconds. If the mouse entered its head into the port while the CS+ was being presented, reward was delivered. A mouse entering its head into the port during CS– presentation was not rewarded. During conditioning, microendoscopic imaging was performed on alternating imaging sessions. The terms naïve, intermediate and expert sessions refer to the 1^st^, 7^th^ (9^th^ in one mouse) and 15^th^ (13^th^ in one mouse) training sessions, respectively. Advanced training sessions are sessions 11, 13 and 15. Once the association between the CS+ and the reward was fully formed, mice underwent an extinction paradigm. Extinction sessions were identical to conditioning sessions, but no reward was delivered. The behavioral paradigm was designed using the Med-PC software suite (Med associates, Fairfax, VT, USA). Training took place during the dark phase of the reversed light-dark cycle.

### Monitoring and analyzing mouse kinematics

In all imaging sessions, mouse kinematics were monitored using an accelerometer (HARP, Champalimaud Foundation Hardware Platform, Lisbon, Portugal) and two video cameras (Firefly MV FFMV-03M2M, FLIR Systems, Portland, Oregon, USA), which were all synchronized with the microendoscope. The accelerometer was mounted on the mouse’s head and recorded the body acceleration in the anterio-posterior (BA_AP_), mediolateral (BA_ML_) and dorsoventral (BA_DV_) axes. As previously described(Klaus et al., 2017), the total body acceleration (BA) was defined as:

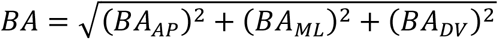

Movement onset and offset times were detected by setting a threshold on the total body acceleration.For each mouse and each session, we plotted the distribution of total body acceleration values. The distributions were bimodal, and the threshold was manually set as the middle point between the two peaks (Fig. S2 A-B). Frames in which the acceleration trace was above the threshold were considered movement frames, and frames in which the acceleration trace was below the threshold were considered rest frames. The average threshold (mean ±SD) was 0.0584 ±0.0076 g (the gravitational acceleration). To verify the reliability of movement onset and offset detection, the results for each session were spot-checked against video recordings.

To identify movement onset events accompanied by small or large acceleration changes, we detected the maximal value of the acceleration in a 2 second time window following each movement onset. We then subtracted from the maximal value the value of the acceleration at the time of movement initiation to obtain the amplitude of acceleration change. The amplitudes were divided into quadrants. Movement onsets corresponding to acceleration changes in the 1^st^ quadrant were considered small acceleration change movements. Movement onsets with acceleration changes in the 4^th^ quadrant were large acceleration change events. The identification of CS+ presentation events accompanied by small or large acceleration changes was conducted similarly.

Ipsi- and contra-lateral turns were detected based on movies of mouse behavior recorded using an over-head video camera. We tracked the position of the mouse by applying the DeepLabCut algorithm (Mathis et al., 2018) to video recordings. Results were then used in combination with a custom-made Matlab (MathWorks, Natick, MA, USA) code to identify times in which the mouse performs right and left turns.

### Immunohistochemistry

Mice were deeply anesthetized and perfused transcardially with 0.1 M phosphate buffer (PB) followed by ice-cold 4% paraformaldehyde. 50 μm coronal sections of the striatum were incubated overnight at 4°C in CAS-block (Life Technologies) with a single primary antibody (see Table 1). On the second day, sections were washed in phosphate buffered saline (PBS) and incubated for 3 hours at room temperature with the appropriate fluorophore-conjugated species-specific secondary antibody: Cy3 donkey anti-goat (1:1000, Abcam), Cy3 donkey anti-mouse (1:1000, Jackson ImmunoResearch Laboratories), Cy3 donkey anti-sheep (1:1000, Jackson ImmunoResearch Laboratories), or Alexa 647 donkey anti-rabbit (1:1000, Abcam). Brain sections were rinsed in PBS and directly cover-slipped by fluorescent mounting medium (VECTASHIELD, Vector Laboratories). Sections were imaged using a laser-scanning confocal microscope (Nikon A1 Plus, Nikon Corporation) using a 20X lens (NA: 0.75).

**Table 1.** Primary antibodies used in this study.

| Molecular marker | Host animal | Dilution | Source and catalog number | B-cell lineage | Research Resource Identifier (RRID) |
| --- | --- | --- | --- | --- | --- |
| Choline acetyltransferase (ChAT) | Goat | 1:100 | Millipore (AB144P) | Polyclonal | AB_90661 |
| DARPP32 | Mouse | 1:50 | Santa Cruz Biotechnology (SC-271111) | Monoclonal | AB_10610055 |
| Parvalbumin (PV) | Sheep | 1:500 | R&D Systems (AF 5058) | Polyclonal | AB_2173907 |
| Somatostatin (SOM) | Rabbit | 1:100 | Peninsula Laboratories (T-4102) | Polyclonal | AB_518613 |

### Data analysis

Data analysis was conducted using custom-made Matlab (MathWorks, Natick, MA, USA) code. Peak events in Ca^2+^ traces were detected using a Matlab peak-finding algorithm with the condition that the peak is larger than 2 standard deviation (SD) of the Ca^2+^ trace, as well as the absolute value of the largest negative deflection in the signal. Peristimulus time histograms (PSTHs) around various behavioral events were generated based on these detected Ca^2+^ peaks.

Event triggered averages (ETAs) were calculated for individual neurons by considering a 5 or 10 second time window around all behavioral events of a given type (e.g., movement onset, cue presentation, reward delivery) and averaging the Ca^2+^ or acceleration trace across events.

To divide significantly responsive neurons into positively and negatively modulated neurons, we calculated the average ETA during the 1 second time windows immediately before and after event onset. If the average after the event onset was larger than the average before it, the neuron was considered positively modulated. Otherwise, it was considered negatively modulated.

To calculate pairwise correlations during rest, we detected rest events that lasted at least 2.5 seconds and considered a 2 second time window starting 250 milliseconds after movement termination. For each pair of neurons, we calculated the correlations between their Ca^2+^ traces for each of these 2 second time windows and then averaged over the various rest events, resulting in a single correlation value for each pair of neurons. Pairwise correlations around movement onset were calculated the same way, but a 2 second time window centered around movement onset was considered for every movement event.

To test whether pairwise correlations depend on the distance between neurons, we first constructed a frequency plot of the distance between neuronal pairs’ centers (bin size 10 μm). Then, we averaged the correlations between neuronal pairs belonging to the same bin, such that each distance bin was represented by a single average correlation value. A Matlab function was then used to fit the data with a linear regression model.

To quantify response reliability around movement onset and CS+ presentation, we considered for each event type only neurons whose signals were significantly positively modulated around the onset of this event. For each neuron and each event type, we measured the portion of the behavioral events for which the neuron produced a significant Ca^2+^ transient in a 1 second window following event onset.

To determine the temporal relationship between somatic and annular signals, we detected peaks in the somatic Ca^2+^ trace and averaged both somatic and annular signals around these times. Signals were first averaged over all of the events in each soma-annulus pair, and the resulting traces were then averaged over the pairs. Decay time constants τwere extracted by fitting a decaying exponent (*a · exp(-t/τ*)+*b*, where *t* is time and *a* and *b* are constants) to the average signals for each soma-annulus pair. For the purpose of this analysis only, somatic traces were not extracted using CNMF-E, as was done in the rest of the manuscript. For each somatic ROI, the raw fluorescence trace of the corresponding annulus was subtracted. Somatic ΔF/F traces were then extracted based on the subtracted traces using the approach described above for annular traces.

### Statistics

The nonparametric two-tailed Wilcoxon rank-sum test (RST) was used for non-matched samples, and the nonparametric Wilcoxon signed-rank test (SRT) was used for matched samples. Shaded areas around curves represent standard errors of the mean.

To determine whether an individual neuron significantly modulated its activity around a given event, a bootstrapping-based strategy was employed (Fig. S2). First, we detected instances in which the neuron’s Ca^2+^ ETA crossed a threshold (the 85^th^ percentile). For each such instance, we counted the number of consecutive frames that the ETA spent above the threshold. We next identified the instance corresponding to the largest number of consecutive frames and calculated the area between the ETA and the horizontal line representing the threshold for that instance (Fig. S2D). To create simulated samples, each neuron’s Ca^2+^ trace was circularly shifted by a random number of frames, and the ETA and area were recalculated around the same event times. This was repeated 2,000 or 5,000 times to generate a null distribution (Fig. S2E).

Outliers were detected using a Matlab outlier detection algorithm. A data point was considered an outlier if its value was more than 1.5 interquartile ranges above the upper quartile (75 percent) or below the lower quartile (25 percent). Null hypotheses were rejected if the P-value was <0.05.

### Joint peristimulus time histogram (JPSTH) calculation

The joint peristimulus time histogram (JPSTH) conveys the temporal dynamics of noise correlation between a pair of simultaneously imaged neurons (Aertsen et al., 1989; Gerstein and Perkel, 1969). Each repetition of the relevant behavioral event (the onset of a spontaneous movement or CS+ presentation) was considered a separate trial. For each pair of neurons, we calculated the raw JPSTH matrix, in which the value in the t_1_-t_2_-th bin represents the number of trials in which neuron 1 and neuron 2 produced Ca^2+^transients in time bins t_1_ and t_2_, respectively.

Cells modulated the rate of their Ca^2+^ transients around the various behavioral events. To account for the correlations due to the co-variation in Ca^2+^ signals, the shift predictor matrix was calculated and subtracted from the raw JPSTH (Aertsen et al., 1989). The calculation of the shift predictor matrix was identical to that of the raw JPSTH, except that neuron 2 trials were circularly shifted. Namely, the *n*^th^ trial in neuron 1 was compared to the (*n+m)^th^* trial in neuron 2, where *m* is the shift. We repeated this for all possible shifts and averaged the resulting matrices to obtain the shift predictor matrix. The corrected JPSTH was then derived by subtracting the shift predictor bin by bin from the raw JPSTH and smoothed using a twodimensional Gaussian window with a 1 bin SD. The term ‘JPSTH’ refers to the corrected JPSTH. To normalize the JPSTH of a pair of neurons, we divided it by the SDs of their PSTHs (Aertsen et al., 1989; Joshua et al., 2009). The population JPSTH is the average of the normalized corrected JPSTHs of all relevant neuronal pairs. All calculations were performed in 67-ms-wide bins. However, the JPSTH is unitless and values do not depend on bin width.

To compare the peaks in the diagonal of the population JPSTH to random fluctuations in JPSTH values, we estimated the mean and SD of the diagonal in a 10 second window far removed (a random shift of least 20 seconds) from the behavioral events.

## Data and code availability

Any data or code reported in this paper, as well as any other information required to reanalyze the data, are available from the lead contact upon request.

## References

Aertsen, A.M.H.J., Gerstein, G.L., Habib, M.K., and Palm, G. (1989). Dynamics of neuronal firing correlation: Modulation of “effective connectivity.” J. Neurophysiol. 61. https://doi.org/10.1152/jn.1989.61.5.900.

Albin, R.L., Young, A.B., and Penney, J.B. (1989). The functional anatomy of basal ganglia disorders. Trends Neurosci. 12, 366–375. https://doi.org/10.1016/0166-2236(89)90074-X.

Alexander, G.E., and Crutcher, M.D. (1990). Functional architecture of basal ganglia circuits: neural substrates of parallel processing. Trends Neurosci. 13, 266–271. https://doi.org/10.1016/0166-2236(90)90107-L.

Aosaki, T., Kimura, M., and Graybiel, A.M. (1995). Temporal and Spatial Characteristics of Tonically Active Neurons of the Primate’s Striatum.

Apicella, P., Legallet, E., and Trouche, E. (1997). Responses of tonically discharging neurons in the monkey striatum to primary rewards delivered during different behavioral states. Exp. Brain Res. 116, 456–466. https://doi.org/10.1007/PL00005773.

Assous, M., and Tepper, J.M. (2018). Excitatory extrinsic afferents to striatal interneurons and interactions with striatal microcircuitry. Eur. J. Neurosci. 49, 593–603. https://doi.org/10.1111/ejn.13881.

Bakhurin, K.I., Mac, V., Golshani, P., and Masmanidis, S.C. (2016). Temporal correlations among functionally specialized striatal neural ensembles in reward-conditioned mice. J. Neurophysiol. 115, 1521–1532. https://doi.org/10.1152/JN.01037.2015.

Barbera, G., Liang, B., Zhang, L., Gerfen, C.R., Culurciello, E., Chen, R., Li, Y., and Lin, D.T. (2016). Spatially Compact Neural Clusters in the Dorsal Striatum Encode Locomotion Relevant Information. Neuron 92, 202–213. https://doi.org/10.1016/J.NEURON.2016.08.037.

Benedetti, B.L., Glazewski, S., and Barth, A.L. (2009). Reliable and Precise Neuronal Firing during Sensory Plasticity in Superficial Layers of Primary Somatosensory Cortex. J. Neurosci. 29, 11817–11827. https://doi.org/10.1523/JNEUROSCI.3431-09.2009.

Bennett, B.D., Callaway, J.C., and Wilson, C.J. (2000). Intrinsic membrane properties underlying spontaneous tonic firing in neostriatal cholinergic interneurons. J. Neurosci. 20, 8493–8503. https://doi.org/https://doi.org/10.1523/JNEUROSCI.20-22-08493.2000.

Berke, J.D. (2011). Functional properties of striatal fast-spiking interneurons. Front. Syst. Neurosci. 0, 45. https://doi.org/10.3389/FNSYS.2011.00045/BIBTEX.

Bolam, J.P., Smith, Y., Ingham, C.A., von Krosigk, M., and Smith, A.D. (1993). Chapter 5 Convergence of synaptic terminals from the striatum and the globus pallidus onto single neurones in the substantia nigra and the entopeduncular nucleus. Prog. Brain Res. 99, 73–88. https://doi.org/10.1016/S0079-6123(08)61339-4.

Brown, P. (2007). Abnormal oscillatory synchronisation in the motor system leads to impaired movement. Curr. Opin. Neurobiol. 17, 656–664. https://doi.org/10.1016/j.conb.2007.12.001.

Bullock, T.H. (1970). The Reliability of Neurons. J. Gen. Physiol. 55, 565–584. https://doi.org/10.1085/JGP.55.5.565.

Chan, S.C., Surmeier, J.D., and Yung, W.H. (2006). Striatal information signaling and integration in globus pallidus: Timing matters. NeuroSignals 14, 281–289. https://doi.org/10.1159/000093043.

Chen, T.W., Wardill, T.J., Sun, Y., Pulver, S.R., Renninger, S.L., Baohan, A., Schreiter, E.R., Kerr, R.A., Orger, M.B., Jayaraman, V., et al. (2013). Ultrasensitive fluorescent proteins for imaging neuronal activity. Nature 499, 295–300. https://doi.org/10.1038/nature12354.

Cui, G., Jun, S.B., Jin, X., Pham, M.D., Vogel, S.S., Lovinger, D.M., and Costa, R.M. (2013). Concurrent activation of striatal direct and indirect pathways during action initiation. Nat. 2013 4947436 494, 238–242. https://doi.org/10.1038/nature11846.

DeLong, M. (1990). Primate models of movement disorders of basal ganglia origin. Trends Neurosci. 13, 281–285. https://doi.org/10.1016/0166-2236(90)90110-V.

Doig, N.M., Moss, J., and Bolam, J.P. (2010). Cortical and Thalamic Innervation of Direct and Indirect Pathway Medium-Sized Spiny Neurons in Mouse Striatum. J. Neurosci. 30, 14610–14618. https://doi.org/10.1523/JNEUROSCI.1623-10.2010.

Flaherty, A.W., and Graybiel, A.M. (1993). Two input systems for body representations in the primate striatal matrix: experimental evidence in the squirrel monkey. J. Neurosci. 13, 1120–1137. https://doi.org/10.1523/JNEUROSCI.13-03-01120.1993.

Fobbs, W.C., Bariselli, S., Licholai, J.A., Miyazaki, N., Matikainen-Ankney, B.A., Creed, M.C., and Kravitz, A. V (2020). Continuous representations of speed by striatal medium spiny neurons. J. Neurosci. 40, 1679–1688. https://doi.org/10.1523/JNEUROSCI.1407-19.2020.

Gallant, J.L., and Vinje, W.E. (2001). Reverse Spikeology: Predicting Single Spikes. Neuron 30, 646–647. https://doi.org/10.1016/S0896-6273(01)00336-1.

Gerfen, C.R. (1988). Synaptic organization of the striatum. J. Electron Microsc. Tech. 10, 265–281. https://doi.org/10.1002/jemt.1060100305.

Gerfen, C., Engber, T., Mahan, L., Susel, Z., Chase, T., Monsma, F., and Sibley, D. (1990). D1 and D2 dopamine receptor-regulated gene expression of striatonigral and striatopallidal neurons. Science 250, 1429–1432. https://doi.org/10.1126/SCIENCE.2147780.

Gerstein, G.L., and Perkel, D.H. (1969). Simultaneously recorded trains of action potentials: Analysis and functional interpretation. Science. 164, 828–830. https://doi.org/10.1126/science.164.3881.828.

Gittis, A.H., Leventhal, D.K., Fensterheim, B.A., Pettibone, J.R., Berke, J.D., and Kreitzer, A.C. (2011). Selective Inhibition of Striatal Fast-Spiking Interneurons Causes Dyskinesias. J. Neurosci. 31, 15727–15731. https://doi.org/10.1523/JNEUROSCI.3875-11.2011.

Graveland, G.A., and Difiglia, M. (1985). The frequency and distribution of medium-sized neurons with indented nuclei in the primate and rodent neostriatum. Brain Res. 327, 307–311. https://doi.org/10.1016/0006-8993(85)91524-0.

Gritton, H.J., Howe, W.M., Romano, M.F., DiFeliceantonio, A.G., Kramer, M.A., Saligrama, V., Bucklin, M.E., Zemel, D., and Han, X. (2019). Unique contributions of parvalbumin and cholinergic interneurons in organizing striatal networks during movement. Nat. Neurosci. 22, 586–597. https://doi.org/10.1038/S41593-019-0341-3.

Gur, M., and Snodderly, D.M. (2006). High Response Reliability of Neurons in Primary Visual Cortex (V1) of Alert, Trained Monkeys. Cereb. Cortex 16, 888–895. https://doi.org/10.1093/CERCOR/BHJ032.

Gutkin, B., Ermentrout, G.B., and Rudolph, M. (2003). Spike Generating Dynamics and the Conditions for Spike-Time Precision in Cortical Neurons. J. Comput. Neurosci. 2003 151 15, 91–103. https://doi.org/10.1023/A:1024426903582.

Hammond, C. (2015). Firing patterns of neurons. Cell. Mol. Neurophysiol. Fourth Ed. 343–360. https://doi.org/10.1016/B978-0-12-397032-9.00017-0.

Hikosaka, O., Takikawa, Y., and Kawagoe, R. (2000). Role of the basal ganglia in the control of purposive saccadic eye movements. Physiol. Rev. 80, 953–978. https://doi.org/10.1152/PHYSREV.2000.80.3.953.

Hunnicutt, B.J., Jongbloets, B.C., Birdsong, W.T., Gertz, K.J., Zhong, H., and Mao, T. (2016). A comprehensive excitatory input map of the striatum reveals novel functional organization. eLife 5. https://doi.org/10.7554/eLife.19103.

Hunter, J.D., and Milton, J.G. (2003). Amplitude and frequency dependence of spike timing: Implications for dynamic regulation. J. Neurophysiol. 90, 387–394. https://doi.org/10.1152/JN.00074.2003/ASSET/IMAGES/LARGE/9K0733181006.JPEG.

Inase, M., Sakai, S.T., and Tanji, J. (1996). Overlapping Corticostriatal Projections From the Supplementary Motor Area and the Primary Motor Cortex in the Macaque Monkey: An Anterograde Double Labeling Study. J. Comp. Neurol. 373283, 296. https://doi.org/10.1002/(SICI)1096-9861(19960916)373:2.

Isomura, Y., Takekawa, T., Harukuni, R., Handa, T., Aizawa, H., Takada, M., and Fukai, T. (2013). Reward-Modulated Motor Information in Identified Striatum Neurons. J. Neurosci. 33, 10209–10220. https://doi.org/10.1523/JNEUROSCI.0381-13.2013.

Jin, X., Tecuapetla, F., and Costa, R.M. (2014). Basal ganglia subcircuits distinctively encode the parsing and concatenation of action sequences. Nat. Neurosci. 2014 173 17, 423–430. https://doi.org/10.1038/nn.3632.

Johansson, Y., and Silberberg, G. (2020). The Functional Organization of Cortical and Thalamic Inputs onto Five Types of Striatal Neurons Is Determined by Source and Target Cell Identities. Cell Rep. 30, 1178–1194.e3. https://doi.org/10.1016/J.CELREP.2019.12.095.

Jones, L.M., Depireux, D.A., Simons, D.J., and Keller, A. (2004). Robust temporal coding in the trigeminal system. Science (80-.). 304, 1986–1989. https://doi.org/10.1126/SCIENCE.1097779/SUPPL_FILE/JONES.SOM.PDF.

Joshua, M., Adler, A., Prut, Y., Vaadia, E., Wickens, J.R., and Bergman, H. (2009). Synchronization of Midbrain Dopaminergic Neurons Is Enhanced by Rewarding Events. Neuron 62, 695–704. https://doi.org/10.1016/j.neuron.2009.04.026.

JW,M. (1996). The basal ganglia: focused selection and inhibition of competing motor programs. Prog. Neurobiol. 50, 381–425. https://doi.org/10.1016/S0301-0082(96)00042-1.

Kara, P., Reinagel, P., and Reid, R.C. (2000). Low Response Variability in Simultaneously Recorded Retinal, Thalamic, and Cortical Neurons. Neuron 27, 635–646. https://doi.org/10.1016/S0896-6273(00)00072-6.

Kimura, M., Rajkowski, J., and Evarts, E. (1984). Tonically discharging putamen neurons exhibit set-dependent responses. Neurobiology 81, 4998–5001. https://doi.org/10.1073/pnas.81.15.4998.

Kincaid, A.E., Zheng, T., and Wilson, C.J. (1998). Connectivity and convergence of single corticostriatal axons. J. Neurosci. 18, 4722–4731. https://doi.org/10.1523/jneurosci.18-12-04722.1998.

Klaus, A., Martins, G.J., Paixao, V.B., Zhou, P., Paninski, L., and Costa, R.M. (2017). The Spatiotemporal Organization of the Striatum Encodes Action Space. Neuron 95, 1171–1180.e7. https://doi.org/10.1016/j.neuron.2017.08.015.

Koos, T., Tepper, J.M., and Wilson, C.J. (2004). Comparison of IPSCs Evoked by Spiny and Fast-Spiking Neurons in the Neostriatum. J. Neurosci. 24, 7916–7922. https://doi.org/10.1523/JNEUROSCI.2163-04.2004.

Koós, T., and Tepper, J.M. (2002). Dual cholinergic control of fast-spiking interneurons in the neostriatum. J. Neurosci. 22, 529–535. https://doi.org/10.1523/jneurosci.22-02-00529.2002.

Kravitz, A. V., Freeze, B.S., Parker, P.R.L., Kay, K., Thwin, M.T., Deisseroth, K., and Kreitzer, A.C. (2010). Regulation of parkinsonian motor behaviours by optogenetic control of basal ganglia circuitry. Nature 466, 622–626. https://doi.org/10.1038/nature09159.

Legaria, A.A., Yang, B., Ahanonu, B., Licholai, J.A., Parker, J.G., and Kravitz, A. V. (2021). Striatal fiber photometry reflects primarily non-somatic activity. BioRxiv 2021.01.20.427525. https://doi.org/10.1101/2021.01.20.427525.

Liberti, W.A., Schmid, T.A., Forli, A., Snyder, M., and Yartsev, M.M. (2022). A stable hippocampal code in freely flying bats. Nature 604, 98–103. https://doi.org/10.1038/s41586-022-04560-0.

Mainen, Z.F., and Seinowski, T.J. (1995). Reliability of spike timing in neocortical neurons. Science 268, 1503–1506. https://doi.org/10.1126/SCIENCE.7770778.

Mathis, A., Mamidanna, P., Cury, K.M., Abe, T., Murthy, V.N., Mathis, M.W., and Bethge, M. (2018). DeepLabCut: markerless pose estimation of user-defined body parts with deep learning. Nat. Neurosci. 21, 1281–1289. https://doi.org/10.1038/s41593-018-0209-y.

McFarland, N.R., and Haber, S.N. (2000). Convergent inputs from thalamic motor nuclei and frontal cortical areas to the dorsal striatum in the primate. J. Neurosci. 20, 3798–3813. https://doi.org/10.1523/JNEUROSCI.20-10-03798.2000.

Nambu, A. (2008). Seven problems on the basal ganglia. Curr. Opin. Neurobiol. 18, 595–604. https://doi.org/10.1016/J.CONB.2008.11.001.

Naundorf, B., Wolf, F., and Volgushev, M. (2006). Unique features of action potential initiation in cortical neurons. Nature 440, 1060–1063. https://doi.org/10.1038/nature04610.

Nolte, M., Reimann, M.W., King, J.G., Markram, H., and Muller, E.B. (2019). Cortical reliability amid noise and chaos. Nat. Commun. 2019 101 10, 1–15. https://doi.org/10.1038/s41467-019-11633-8.

Ouimet, C.C., Langley-Gullion, K.C., and Greengard, P. (1998). Quantitative immunocytochemistry of DARPP-32-expressing neurons in the rat caudatoputamen. Brain Res. 808, 8–12. https://doi.org/10.1016/S0006-8993(98)00724-0.

Parker, J.G., Marshall, J.D., Ahanonu, B., Wu, Y.W., Kim, T.H., Grewe, B.F., Zhang, Y., Li, J.Z., Ding, J.B., Ehlers, M.D., et al. (2018). Diametric neural ensemble dynamics in parkinsonian and dyskinetic states. Nature 557, 177–182. https://doi.org/10.1038/S41586-018-0090-6.

Percheron, G., Yelnik, J., and François, C. (1984). A Golgi analysis of the primate globus pallidus. III. Spatial organization of the striato-pallidal complex. J. Comp. Neurol. 227, 214–227. https://doi.org/10.1002/cne.902270207.

Percheron, G., François, C., and Yelnik, J. (1987). Spatial Organization and Information Processing in the Core of the Basal Ganglia. (Springer, Boston, MA), pp. 205–226.

Percheron, G., François, C., Yelnik, J., Fénelon, G., and Talbi, B. (1994). The Basal Ganglia Related System of Primates: Definition, Description and Informational Analysis. 3–20. https://doi.org/10.1007/978-1-4613-0485-2_1.

Perk, C.G., Wickens, J.R., and Hyland, B.I. (2015). Differing properties of putative fastspiking interneurons in the striatum of two rat strains. Neuroscience 294, 215–226. https://doi.org/10.1016/J.NEUROSCIENCE.2015.02.051.

Plotkin, J.L., Day, M., and Surmeier, D.J. (2011). Synaptically driven state transitions in distal dendrites of striatal spiny neurons. Nat. Neurosci. 14, 881–888. https://doi.org/10.1038/nn.2848.

Pnevmatikakis, E.A., Soudry, D., Gao, Y., Machado, T.A., Merel, J., Pfau, D., Reardon, T., Mu, Y., Lacefield, C., Yang, W., et al. (2016). Simultaneous Denoising, Deconvolution, and Demixing of Calcium Imaging Data. Neuron 89, 299. https://doi.org/10.1016/j.neuron.2015.11.037.

Puelma Touzel, M., and Wolf, F. (2015). Complete Firing-Rate Response of Neurons with Complex Intrinsic Dynamics. PLOS Comput. Biol. 11, e1004636. https://doi.org/10.1371/journal.pcbi.1004636.

Raz, A., Feingold, A., Zelanskaya, V., Vaadia, E., and Bergman, H. (1996). Neuronal synchronization of tonically active neurons in the striatum of normal and parkinsonian primates. J. Neurophysiol. 76.

Rehani, R., Atamna, Y., Tiroshi, L., Chiu, W.-H., de Jesús Aceves Buendía, J., Martins, G.J., Jacobson, G.A., and Goldberg, J.A. (2019). Activity Patterns in the Neuropil of Striatal Cholinergic Interneurons in Freely Moving Mice Represent Their Collective Spiking Dynamics. Eneuro 6, ENEURO.0351-18.2018. https://doi.org/10.1523/ENEURO.0351-18.2018.

Schreiber, S., Samengo, I., and Herz, A.V.M. (2009). Two distinct mechanisms shape the reliability of neural responses. J. Neurophysiol. 101, 2239–2251. https://doi.org/10.1152/jn.90711.2008.

Shadlen, M.N., and Newsome, W.T. (1998). The Variable Discharge of Cortical Neurons: Implications for Connectivity, Computation, and Information Coding. J. Neurosci. 18, 3870–3896. https://doi.org/10.1523/JNEUROSCI.18-10-03870.1998.

Sheintuch, L., Rubin, A., Brande-Eilat, N., Geva, N., Sadeh, N., Pinchasof, O., and Correspondence, Y.Z. (2017). Tracking the Same Neurons across Multiple Days in Ca2+ Imaging Data. CellReports 21, 1102–1115. https://doi.org/10.1016/j.celrep.2017.10.013.

Shin, J.H., Kim, D., and Jung, M.W. (2018). Differential coding of reward and movement information in the dorsomedial striatal direct and indirect pathways. Nat. Commun. 2018 91 9, 1–14. https://doi.org/10.1038/s41467-017-02817-1.

Shin, J.H., Song, M., Paik, S.B., and Jung, M.W. (2020). Spatial organization of functional clusters representing reward and movement information in the striatal direct and indirect pathways. Proc. Natl. Acad. Sci. U. S. A. 117, 27004–27015. https://doi.org/10.1073/pnas.2010361117.

Slater, C.R. (2008). Reliability of neuromuscular transmission and how it is maintained. Handb. Clin. Neurol. 91, 27–101. https://doi.org/10.1016/S0072-9752(07)01502-3.

Smith, Y., Raju, D. V., Pare, J.F., and Sidibe, M. (2004). The thalamostriatal system: a highly specific network of the basal ganglia circuitry. Trends Neurosci. 27, 520–527. https://doi.org/10.1016/J.TINS.2004.07.004.

Takada, M., Tokuno, H., Nambu, A., and Inase, M. (1998). Corticostriatal input zones from the supplementary motor area overlap those from the contra-rather than ipsilateral primary motor cortex. Brain Res. 791, 335–340. https://doi.org/10.1016/S0006-8993(98)00198-X.

Takadá, M., Tokunó, H., Nambú, A., Inase, M., Takada, M., Tokuno, ’ H, Nambu, A., and Inase, ’ M (1998). Corticostriatal projections from the somatic motor areas of the frontal cortex in the macaque monkey: segregation versus overlap of input zones from the primary motor cortex, the supplementary motor area, and the premotor cortex. Exp. Brain Res. 1998 1201 120, 114–128. https://doi.org/10.1007/S002210050384.

Tchumatchenko, T., Malyshev, A., Wolf, F., and Volgushev, M. (2011). Ultrafast population encoding by cortical neurons. J. Neurosci. 31, 12171–12179.https://doi.org/10.1523/JNEUROSCI.2182-11.2011.

Tecuapetla, F., Matias, S., Dugue, G.P., Mainen, Z.F., and Costa, R.M. (2014). Balanced activity in basal ganglia projection pathways is critical for contraversive movements. Nat. Commun. 2014 51 5, 1–10. https://doi.org/10.1038/ncomms5315.

Tepper, J.M. (2010). GABAergic Interneurons of the Striatum. In Handbook of Basal Ganglia Structure and Function, (Elsevier), pp. 151–166.

Tepper, J.M., and Koós, T. (2016). GABAergic Interneurons of the Striatum. Handb. Behav. Neurosci. 24, 157–178. https://doi.org/10.1016/B978-0-12-802206-1.00008-8.

Tepper, J.M., Wilson, C.J., and Koós, T. (2008). Feedforward and feedback inhibition in neostriatal GABAergic spiny neurons. Brain Res. Rev. 58, 272–281. https://doi.org/10.1016/j.brainresrev.2007.10.008.

Yeterian, E.H., and Van Hoesen, G.W. (1978). Cortico-striate projections in the rhesus monkey: The organization of certain cortico-caudate connections. Brain Res. 139, 43–63. https://doi.org/10.1016/0006-8993(78)90059-8.

Zheng, T., and Wilson, C.J. (2002). Corticostriatal combinatorics: The implications of corticostriatal axonal arborizations. J. Neurophysiol. 87, 1007–1017. https://doi.org/10.1152/jn.00519.2001.

Zhou, P., Resendez, S.L., Rodriguez-Romaguera, J., Jimenez, J.C., Neufeld, S.Q., Stuber, G.D., Hen, R., Kheirbek, M.A., Sabatini, B.L., Kass, R.E., et al. (2018). Efficient and accurate extraction of in vivo calcium signals from microendoscopic video data. Elife 7. https://doi.org/10.7554/eLife.28728.

