## Supplemental Figures for "Striatal neurons are recruited dynamically into collective representations of self-initiated and learned actions in freely-moving mice"

### Supplementary information

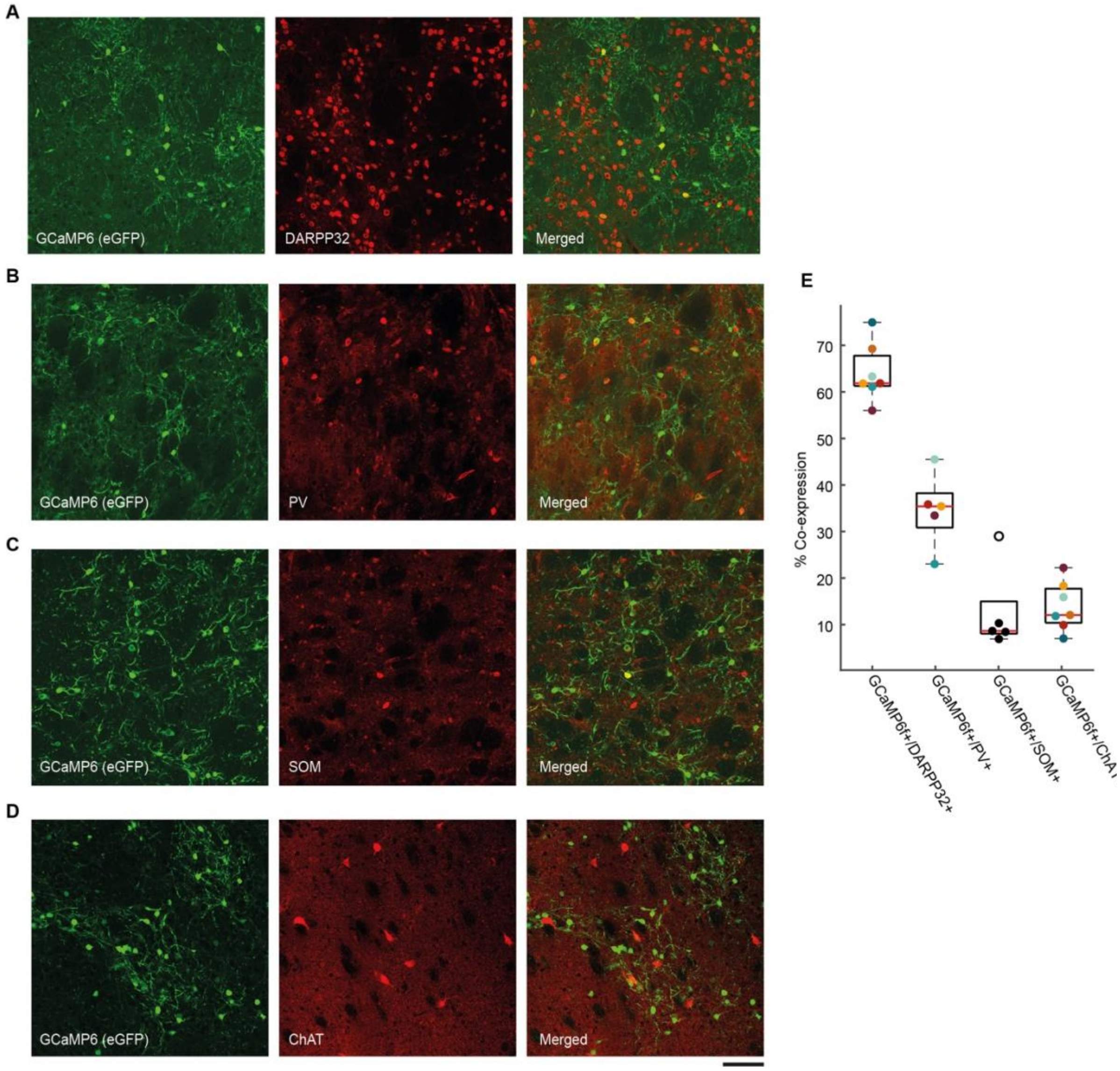


**Figure S1. Striatal neurons in the sparse GCaMP mouse are a mixture of neuronal types with an SPN majority. A:** Immunohistochemical analysis of dorsal striatum of sparse GCaMP mice demonstrates that a large portion of GCaMP6f expressing cells express DARPP32. **B:** Same as A for the co-expression of GCaMP6f and parvalbumin (PV). **C:** Same as A for the co-expression of GCaMP6f and somatostatin (SOM). **D:** Same as A for the co-expression of GCaMP6f and ChAT. Scale bar = 100 μm (A-D). **E:** Rates of co-expression of GCaMP6f and markers for various neuronal subtypes in the striatum. 64±2.3% (mean ± S.E.M; range 56-74.9%, N=7 mice) of GCaMP6f-expressing neurons co-express DARPP32, an established SPN marker(Ouimet et al., 1998); 34.64±3.58% (mean ± S.E.M; range 23-45.5%, N=5 mice) co-express PV; and 8.6±0.7% [mean ± S.E.M; range 6.9-10.3%, N=4 mice; one outlier (29%) was excluded from the analysis] co-express SOM;13.9±2% (mean ± S.E.M; range 7-22.2%, N=7 mice) of GCaMP6f-expressing cells exhibited immunoreactivity to ChAT; Importantly, the analysis for the co-expression of SOM and GCaMP6f was conducted on sparse GCaMP mice that were not included in the microendoscopic experiments. Each data point represents the co-expression rate for a single mouse. Data points with the same color are from the same individual mouse. Black points represent data from sparse GCaMP mice that were not included in the microendoscopic experiments. Outliers are marked by empty circles. Red line is the median. Box edges are 25$^{th}$ and 75$^{th}$ percentile. Whiskers extend to the most extreme data points not considered outliers.

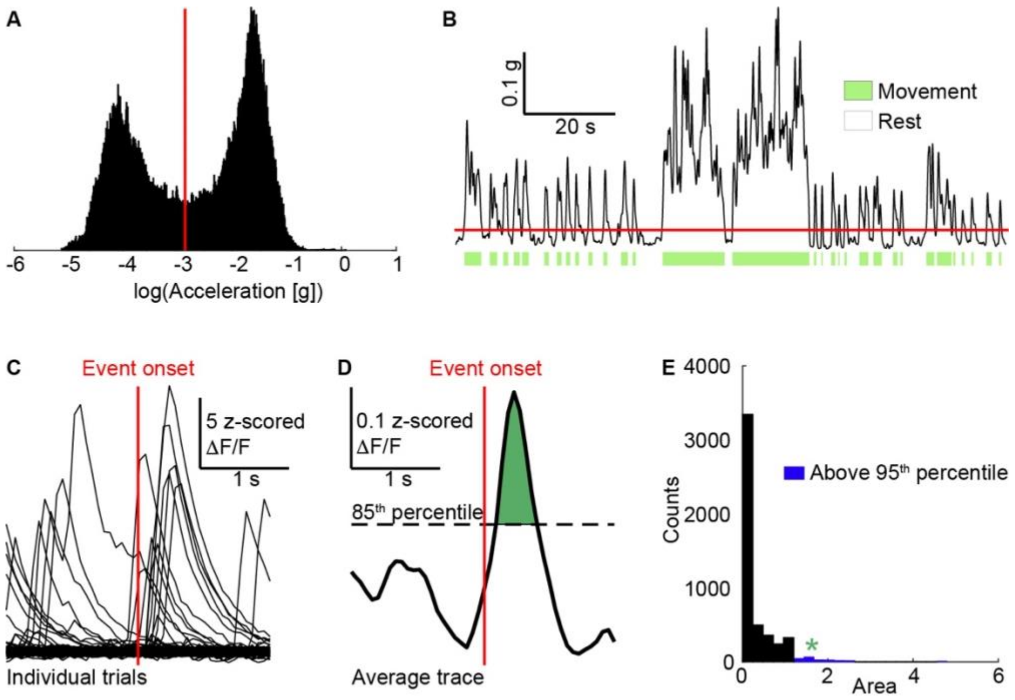


**Figure S2. Detection of movement onset and offset times and determination of response significance A:** Distribution of total body acceleration values from a single session in a representative mouse. Red line represents the threshold, manually set as the middle point between the two peaks in the bimodal distribution. **B:** Total body acceleration trace for the same session. Red line represents the threshold. Green bars mark movement times. **C:** $Ca^{2+}$ activity of a representative neuron around a behavioral event for various trials. Red line marks event onset. **D:** Average response of the same neuron. Dashed line represents an 85$^{th}$ percentile threshold. The statistic for significance testing is the area between the average trace and the threshold (green). **E:** Null distribution of area values generated by bootstrapping. The area for which null-hypothesis is rejected is in blue. The empirical area value for the given neuron is marked by the green asterisk.

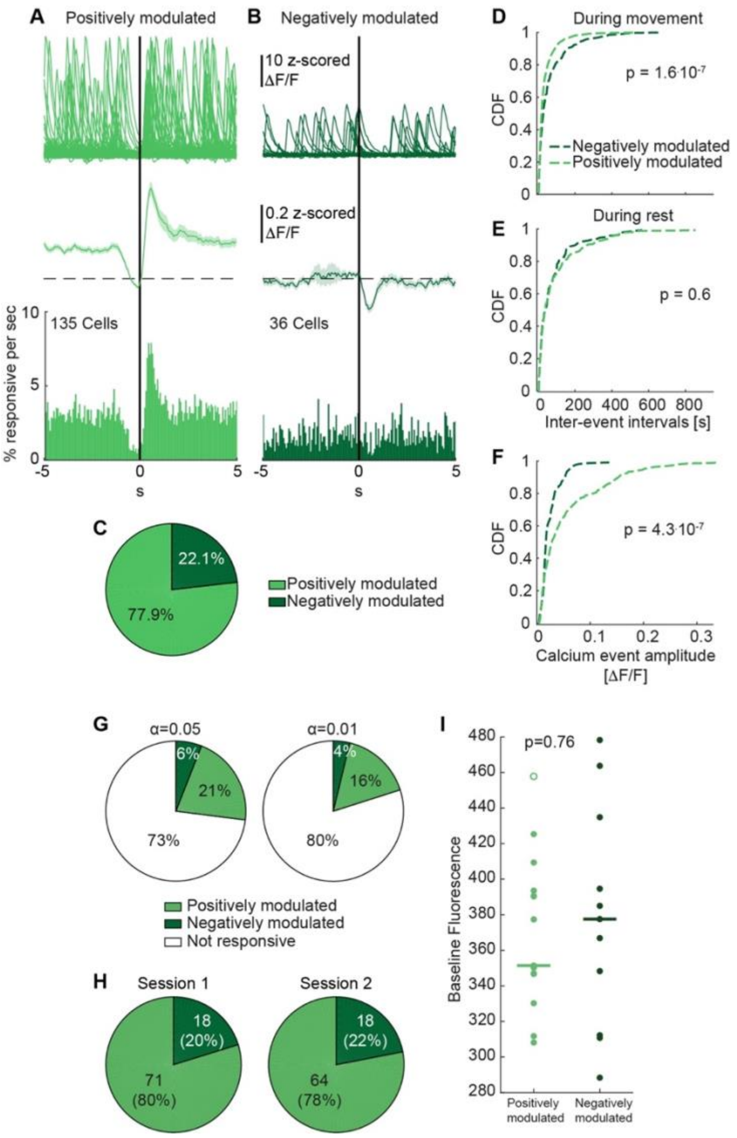


**Figure S3. Two types of responses around self-initiated movement. A:** $Ca^{2+}$ activity around movement onset of a representative positively modulated neuron (top). Each trace represents the activity around a single movement initiation event. The average $Ca^{2+}$ activity across the population of positively modulated neurons (middle). Shaded areas represent S.E.M. PSTH of $Ca^{2+}$ events around movement onset in the population of positively modulated neurons (bottom). **B:** Same as A, for neurons that are negatively modulated around movement onset. **C:** The percentage of positively (light green) and negatively (dark green) modulated neurons, out of the population of significantly modulated neurons. **D:** The CDFs of inter-event intervals for positively (light green) and negatively (dark green) modulated neurons during movement. **E:** Same as D, during rest. **F:** CDFs of $Ca^{2+}$ event amplitudes in positively (light green) and negatively (dark green) modulated neurons during rest. **G:** Rates of imaged neurons positively and negatively modulating their $Ca^{2+}$ signals following movement onset for different significance levels. **H:** Rates of neurons positively and negatively modulating their $Ca^{2+}$ signals following movement onset on the first (left) and second (right) free movement imaging sessions. **I:** Baseline fluorescence values for positively and negatively modulated neurons. Each data point is the average value across all neurons of the relevant category in the given mouse and session. Empty circles represent outliers ($p = 0.76$, SRT).

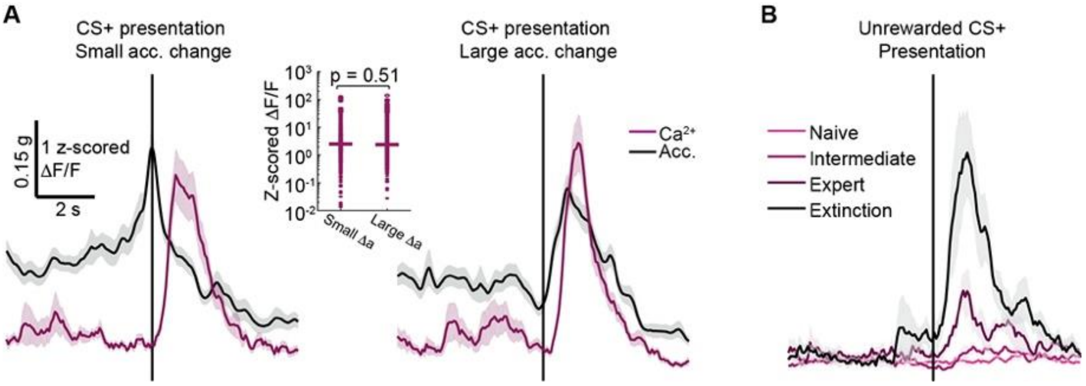


**Figure S4. Responses to cue presentation are not determined by total body acceleration or reward delivery. A:** Average $Ca^{2+}$ activity across the neuronal population (pink) and total body acceleration (black) around CS+ presentations with small (left, 1$^{st}$ quadrant) and large (right, 4$^{th}$ quadrant) acceleration changes. Inset shows distributions of $Ca^{2+}$ response amplitudes for small and large acceleration changes (p=0.51, SRT). **B:** Average $Ca^{2+}$ activity across the population for unrewarded CS+ presentations on various stages of conditioning (light to dark pink), and for the first 10 CS+ presentations on the first extinction session (black). Shaded areas represent S.E.M.

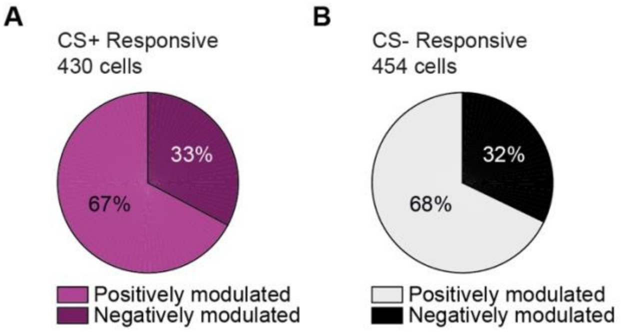


**Figure S5. Striatal neurons exhibit positively and negatively modulated responses around cue presentation in an operant learning paradigm. A:** Percentage of positively (light) and negatively (dark) modulated neurons of CS+ responsive neurons. B**:** Same as F, for CS– presentation.

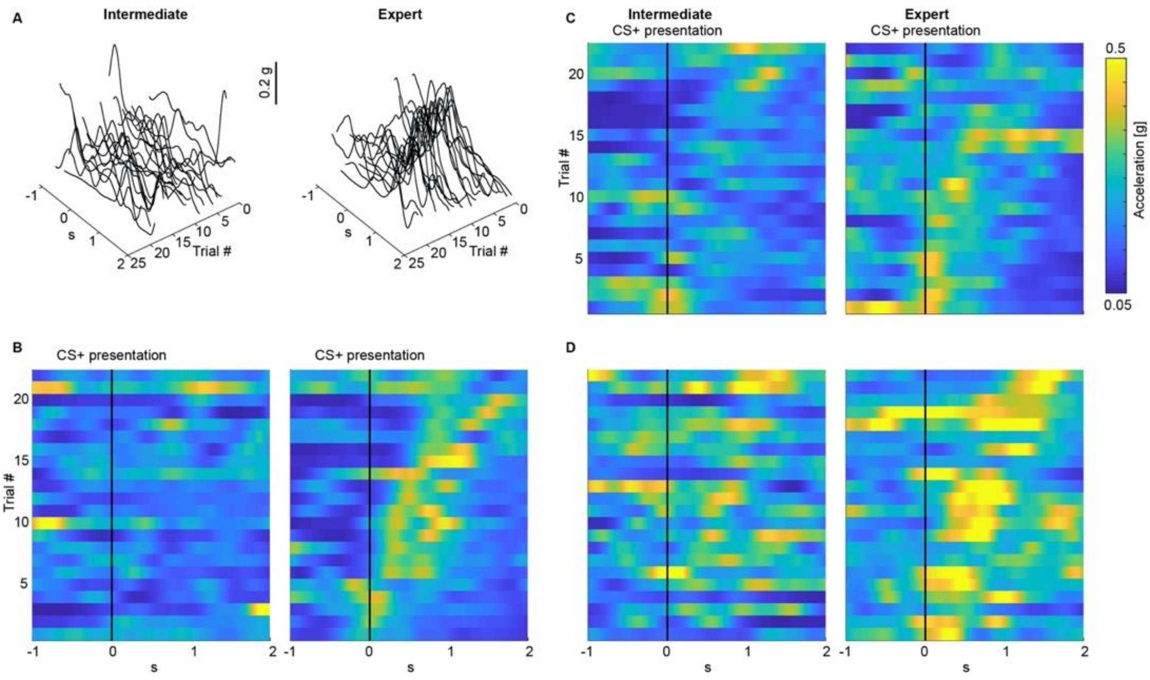


**Figure S6. Mice acceleration profiles following CS+ presentation become more stereotyped as training progresses. A:** Total body acceleration traces from a representative mouse around CS+ presentation on intermediate (left) and expert (right) training sessions. Each trace represents a single trial. **B:** Color-coded matrix showing the total-body acceleration of the same mouse around CS+ presentation on intermediate (left) and expert (right) training sessions. Each row represents a single trial. **C:** Same as B, for a second mouse. **D:** Same as B, for a third mouse.

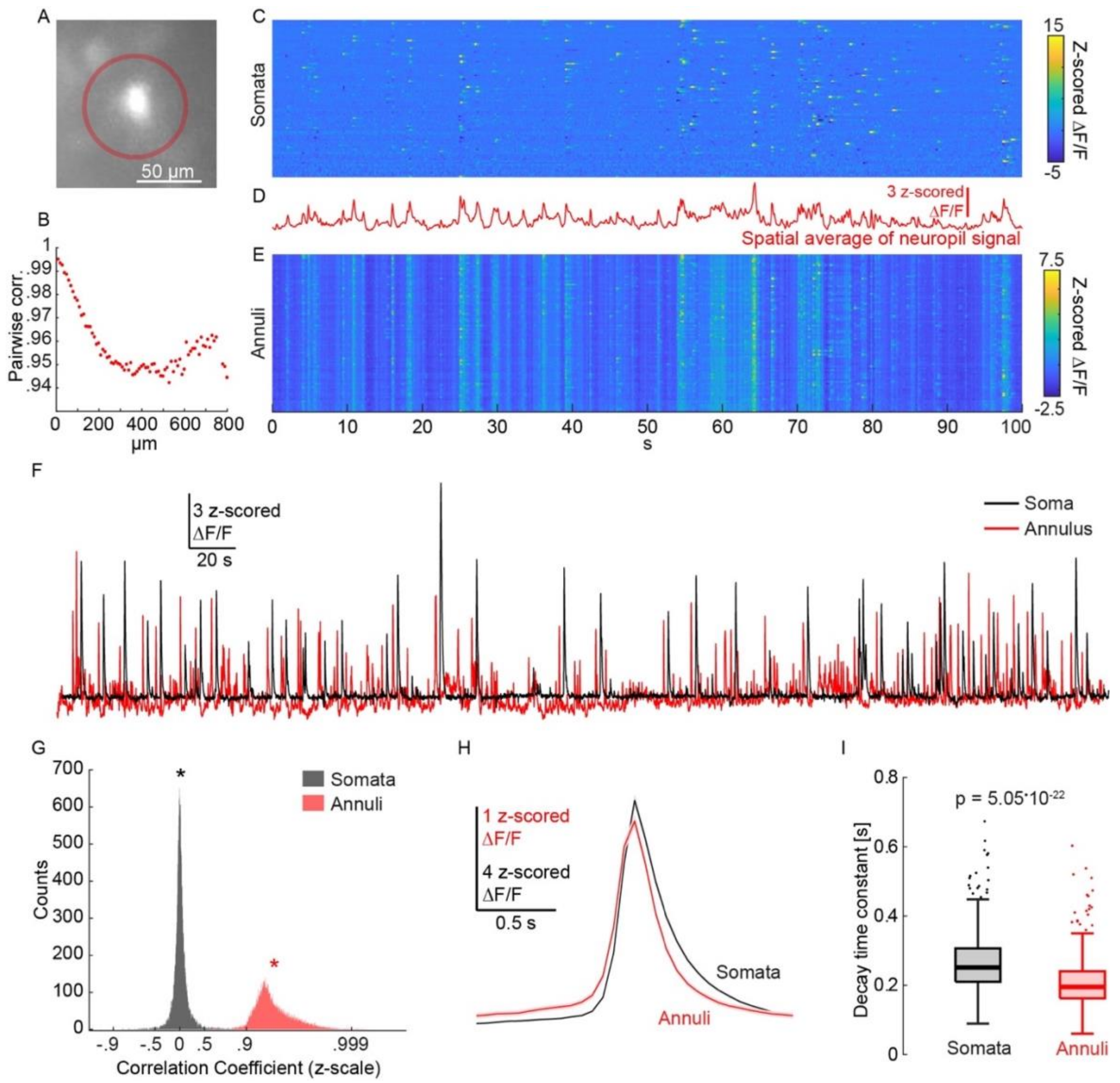


**Figure S7: Neuropil signal is highly correlated in space and displays different kinetics then the somatic signals. A.** Illustration of the sampling of a soma and the surrounding annular region of interest (red). **B.** Pairwise correlations between annular signals as a function of the distance between neuronal centers for all pairs of co-imaged neurons. Each point is the average correlation across all neuronal pairs belonging to the relevant 10 μm distance bin. **C.** Color-coded matrix of the fluctuations in fluorescence as a function of time in 80 somata detected in a single free movement session in a single mouse, with each row representing an individual soma**. D.** Spatial average of signals from all annuli presented in D. **E.** Same as B, with each row representing the signal of the corresponding annulus. **F.** $Ca^{2+}$ signals from a soma-annulus pair. **G.** Distribution of pairwise correlation coefficients for 21,427 simultaneously imaged soma-soma and annulus-annulus pairs. Stars mark the means (0.023 and 0.96 for somata and annuli, respectively). **I.** Average $Ca^{2+}$ signal from the soma and its corresponding annulus averaged over 429 soma-annulus pairs triggered on the somatic $Ca^{2+}$ events. Shaded areas mark S.E.M. **H.** Boxplot of decay time constants for somatic and annular $Ca^{2+}$ signals. The bold like is the median and the whiskers are the 25$^{th}$ and 75$^{th}$ percentiles. Dots represent outliers.
